# Pleiotropic regulation of bacterial toxin production and Allee effect govern microbial predator–prey interactions

**DOI:** 10.1101/2024.07.10.602915

**Authors:** Harikumar R. Suma, Pierre Stallforth

## Abstract

Bacteria are social organisms, which are constantly exposed to predation by nematodes or amoebae. To counteract these predation pressures, bacteria have evolved a variety of potent antipredator strategies. Bacteria of the genus *Pseudomonas*, for instance, evade amoebal predation by the secretion of amoebicidal natural products. The soil bacterium *Pseudomonas fluorescens* HKI0770 produces pyreudione alkaloids that can kill amoebae. Even though the mode of action of the pyreudiones has been elucidated, the spatiotemporal dynamics underlying this predator–prey interaction remain unknown. Using a combination of microscopic and analytical techniques, we elucidated the intricate relationship of this predator−prey association. We used the chromatic bacteria toolbox for intraspecific differentiation of the amoebicide-producing wildtype and the non-producing mutant within microcosms. These allow for variations in nutrient availability and the emergence of predation-evasion strategies of interacting microorganisms. Imaging of the co-cultures revealed that the amoebae initially ingest both the non-producer as well as the toxin-producer cells. The outcomes of predator–prey interactions are governed by the population size and fitness of the interacting partners. We identified that changes in the cell density coupled with alterations in nutrient availability led to a strong Allee effect resulting in the diminished production of pyreudione A. The loss of defence capabilities renders *P. fluorescens* HKI0770 palatable to amoebae. Such a multifaceted regulation provides the basis for a model by which predator–prey populations are being regulated in specific niches. Our results demonstrate how the spatiotemporal regulation of bacterial toxin production alters the feeding behaviour of amoebae.

## Introduction

Bacteria engage in a wide range of social behaviours when interacting with other microorganisms of the same or different species.(Crespi, 2001) Soil-dwelling bacteria for instance are exposed to different bacterivorous microfauna. Grazing by protists in particular regulates bacterial populations and is crucial in shaping natural microbial communities.(Dunn et al., 2018; Jousset, 2012; Klapper et al., 2016; Rossine et al., 2022) From an ecological standpoint, bacterial predation plays a critical role in maintaining the equilibrium of biomes as it curtails the uncontrolled proliferation of certain bacterial species, promoting microbial diversity and nutrient cycling.

Faced with strong predation pressure, bacteria have developed intricate defence mechanisms based on either the creation of physical barriers such as biofilms,(Hahn and Höfle, 2001; Matz and Kjelleberg, 2005) increased motility (Matz and Kjelleberg, 2005) alongside intracellular survival of bacteria within the predator.(Ma et al., 2022; Shi et al., 2021; Strassmann and Shu, 2017) The biosynthesis of toxic low molecular weight natural products, on the other hand, represents a potent defence mechanism to kill adversaries.(Arp et al., 2018; Jousset et al., 2006; Klapper et al., 2016; Matz et al., 2004; Pflanze et al., 2023) The co-evolution of polymicrobial associations has even led to the emergence of cooperative antipredator defence strategies.(Scherlach et al., 2012; Zhang et al., 2021) To counteract these strategies, predators such as amoebae have also evolved efficient strategies to face these mechanisms.(Dinh et al., 2018; Dunn et al., 2018; Farinholt et al., 2019; Iqbal et al., 2014; Zanditenas et al., 2023) Thus, these evolutionary arms races have contributed to the development of bacterial virulence and diverse cell-autonomous defence mechanisms. As a consequence, these microbial predator–prey interactions are important sources of highly adapted natural products with great structural and functional diversity that benefit clinical and agricultural applications.(Götze et al., 2023; Hug et al., 2020; Rutledge and Challis, 2015)

Predation on bacteria can be studied conveniently in the laboratory using the model organism *Dictyostelium discoideum*, a ubiquitous social amoeba. In the vegetative form, the amoeboid cells are motile and can feed on bacteria by phagocytosis. Upon starvation, the individual cells aggregate, generating a fruiting body consisting of a basal plate, a stalk, and a spore-containing sorus on top.(Günther et al., 2022; Schaap, 2011) *D. discoideum* is an excellent platform for studying a wide range of eukaryote–bacteria interactions as it exhibits an intricate relationship with their interacting partners, spanning the whole spectrum of symbiosis – from mutualism to antagonism.(Adiba et al., 2010; Bozzaro and Eichinger, 2011; Shi et al., 2021) A key factor in determining the dynamics of predation is the relationship between population density and fitness of both predator and prey.(MacNulty et al., 2014; Mukherjee and Heithaus, 2013; Schrader and Travis, 2012; Sillo et al., 2011) Studies have shown that the outcome of amoeba−bacteria interactions is strongly mediated by cell density and nutrient availability.(Adiba et al., 2010; Sillo et al., 2011). Such population-dependant outcomes between amoebae and bacteria are known as the Allee effect, where a higher population size is associated with increased individual fitness.(Allee, 1949; DiSalvo et al., 2014; Rubin et al., 2019; Stephens et al., 1999) Allee effects can also occur as a result of social interactions among microorganisms, in particular, a strong Allee effect arises when the population is below a certain threshold. For the social microbe, *D. discoideum* the Allee effect can be observed both during the vegetative and aggregation stages of its life cycle.(Hashimoto et al., 1975; Segota et al., 2022)

*Pseudomonas fluorescens* HKI0770 is a soil bacterium known to secrete extracellular amoebicidal natural products – pyreudiones (pys) – enabling the bacteria to evade predation.(Klapper et al., 2016) However, a mutant unable to produce pyreudiones is palatable to amoeba. This emphasizes the role of natural products in shaping predator−prey interactions. Although the mode of action of the pyreudiones has been shown to rely on the ability to act as a protonophore (Klapper et al., 2019), the dynamics of this predator–prey relationship remain unknown. The constitutive expression of pyreudiones by the bacteria makes this system an excellent model for studying antagonistic microbial predator–prey interactions since amoebal predation is evaded by the bacterium’s production of the amoebicide pyreudione. The mutant strain unable to produce this compound becomes edible highlighting the one-to-one correspondence of pyreudione production and predator evasion.

Here, we examined the predator−prey interaction between *D. discoideum* and, *P. fluorescens* HKI0770 in detail. We visualized the interactions between the two organisms by equipping the amoebicide producer and the non-producer with different fluorescent labels. Intriguingly, time-dependent co-culture studies between amoebae and the fluorescent bacterial strains revealed that amoeba−bacteria interactions are mediated by pleiotropic regulation of toxic small molecules resulting from changes in nutrient availability and a cell density-dependent strong Allee effect, inherent to virtually all ecological niches.

## Results

### Fluorescent labels allow the differentiation of *P. fluorescens* HKI0770 strains

In order to enable the differentiation of *Pseudomonas fluorescens* HKI0770 wildtype (wt) and the *Δpys* mutant, deficient in the production of amoebicidal pyreudiones, we used a differential labelling approach. Both the bacterial strains were labelled using the chromatic bacteria toolbox.(Schlechter et al., 2018) To this end, we chromosomally inserted different fluorescent tags into the genome of each bacterial strain. Specifically, *P. fluorescens* HKI0770 was labelled with mScarlet-I (wt::MRE-145) and *P. fluorescens* HKI0770 Δ*pys* was labelled with mTagBFP2 (Δ*pys*::MRE-140).

For the discrimination of these labelled strains within co-cultures, fluorescence microscopy was performed across different ratios (1:1, 1:3, 3:1) of the individual strains. The differentially tagged cell populations could be unambiguously distinguished using this approach (Figure 1A). Fluorescence emitted by both bacteria was detected with single-cell resolution facilitating the differentiation even within mixtures. This further helped in the quantification of bacterial cell numbers from different initial cell ratios. The percentage of bacteria remained proportional to the initial inoculation ratios (Figure 1B). These ratios are used in subsequent co-culture experiments with amoebae.

**Figure 1.**
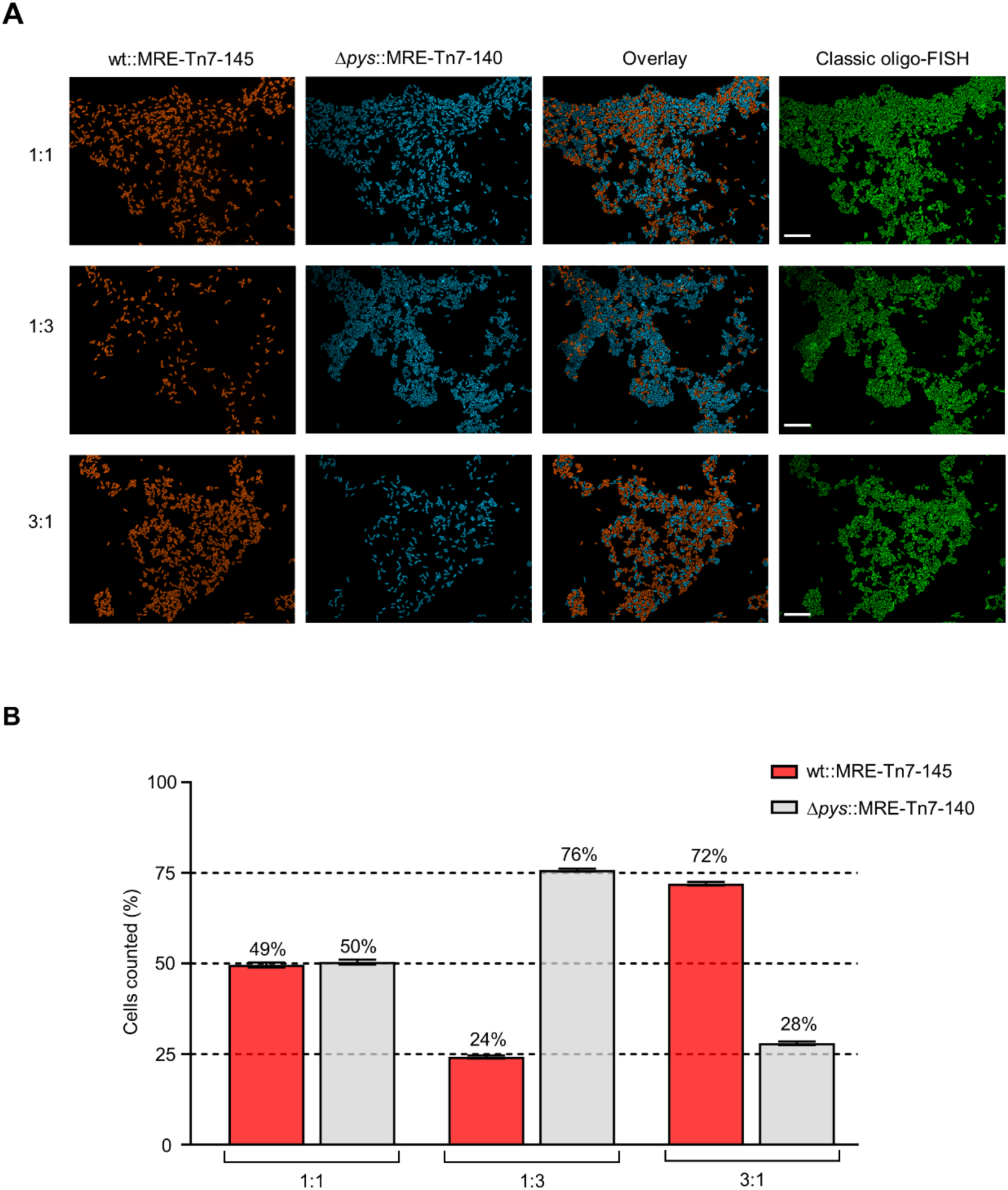
Discrimination between *Pseudomonas fluorescens* HKI0770 strains using fluorescence imaging. **(A)** Representative fluorescence microscopy images of *P. fluorescens* wt::MRE-145 expressing mScarlet-I (red) and *P. fluorescens* Δ*pys*::MRE-140 expressing mTagBFP2 (blue). Different ratios (1:1, 1:3, 3:1) of both strains were used for differentiation. The corresponding FISH image (green) using the PSE227 probe is on the right side. **(B)** Cell counting using CellProfiler software with different ratios of *P. fluorescens* HKI0770 chromatic mutants. Data shown are mean ± standard error (n=10 images per ratio). Graphs are representative of three independent experiments. Black dotted lines represent the reference values (25%, 50%, 75%) for the percentage of cells counted in each ratio. Scale bars are 10 μm. The following source data and figure supplement are available for figure 1: **Source data 1.** Related to Figure 1B. **Figure supplement 1.** The chromatic mutants display same phenotypes as their parent strains.

Importantly, the phenotypes of the chromatic mutants, such as the production of pyreudiones and the feeding behaviour of amoeba, remained unaffected by the labelling (Figure 1—figure supplement 1A and 1B).

### Amoeba can ingest the pyreudione−producing bacteria

The secretion of the amoebicidal natural product, pyreudione A by *P. fluorescens* HKI0770 is an effective strategy to evade predation.(Klapper et al., 2016) However, it remains unknown whether the predator can ingest the toxin producers when pyreudiones are absent in the co-culture.

We prepared co-cultures of *P. fluorescens* HKI0770 chromatic mutants with *D. discoideum* cells expressing GFP-fused phagosomes (See <u>Materials and methods</u>). The co-cultures consisted of bipartite and tripartite combinations of individual chromatic mutants with amoebae.

Live-cell imaging of the co-cultures revealed that *P. fluorescens* wt was indeed ingested by *D. discoideum* cells. The bacteria were observed to be engulfed within the fluorescent phagosomes of the amoebae (Figure 2A). *P. fluorescens Δpys*, which has been already shown to be palatable to amoebae was used as an edible control (Figure 2B). The efficacy of phagocytosis was determined by calculating the phagocytic index (PI) of amoebae. Phagocytic index provides a quantitative measure of phagocytic activity, accounting for both the average number of particles engulfed per amoebae and the ratio of amoebae performing phagocytosis.(Chen et al., 2015; Sano et al., 2003; Santulli-Marotto et al., 2015) A higher PI indicates higher phagocytic activity by phagocytes like amoebae. In the bipartite co-culture, no significant differences between the PIs of *P. fluorescens* wt and Δ*pys* were observed (Figure 2D). This indicates that the amoeba ingested both the amoebicide producer and the non-producer.

**Figure 2.**
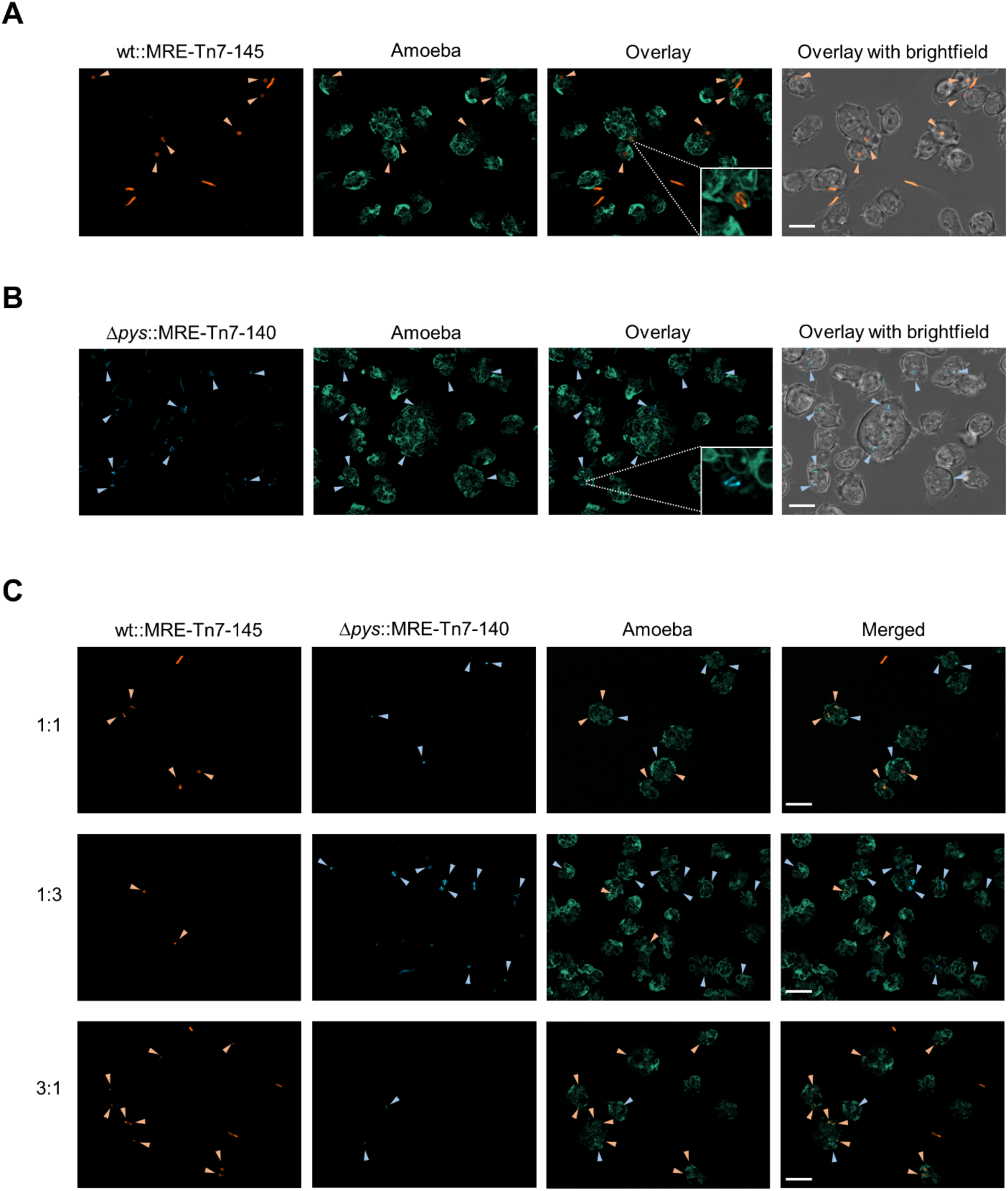
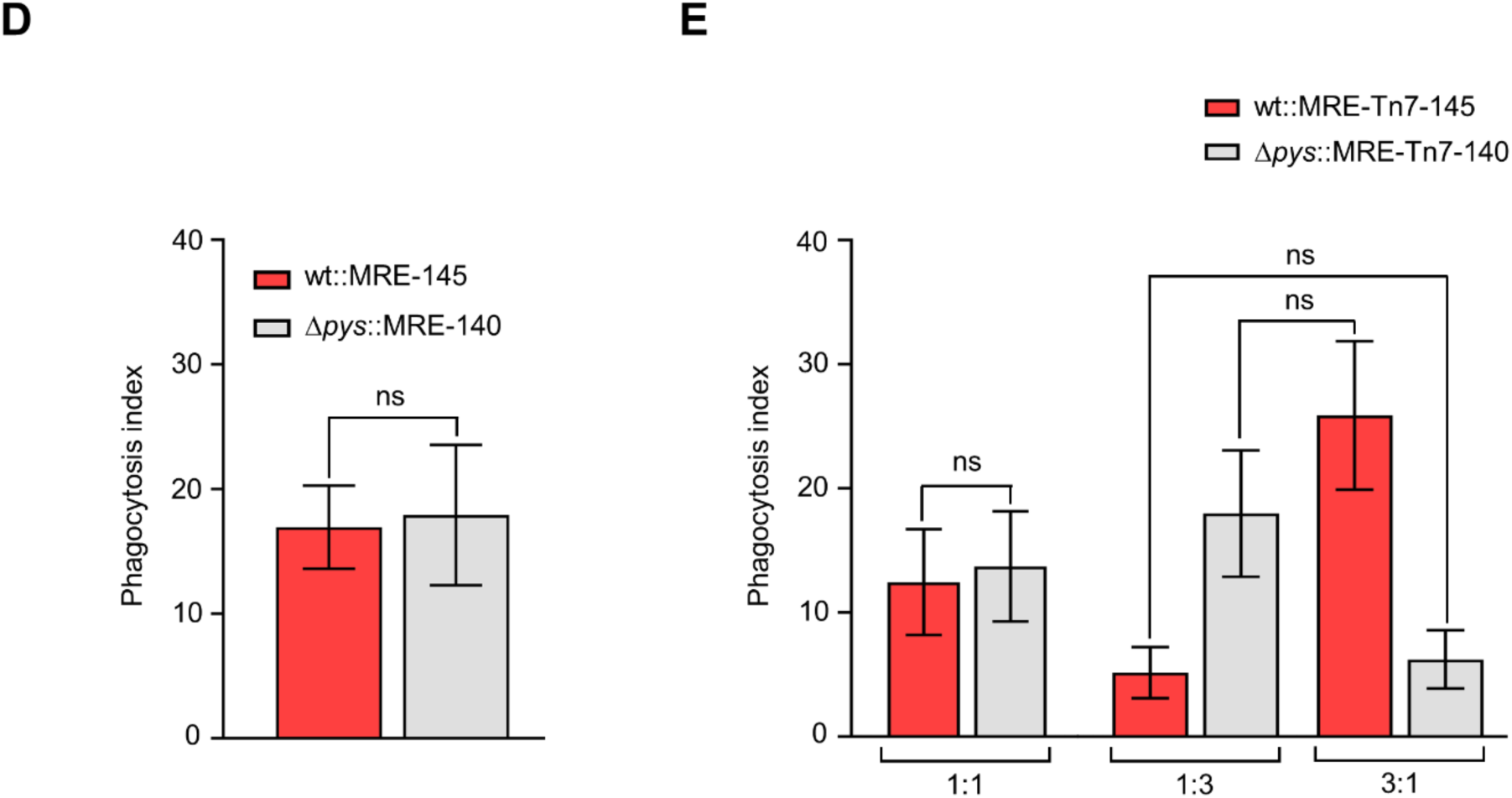
Phagocytosis of *Pseudomonas fluorescens* HKI0770 strains by amoebae. **(A)** A representative 2D maximum projection of the bipartite co-culture between the amoeba, *D. discoideum* vatM:GFP and *P. fluorescens* wt::MRE-145 (orange). **(B)** A representative 2D maximum projection of the bipartite co-culture of amoeba and *P. fluorescens* Δ*pys*::MRE-140 (blue). The insets show engulfed bacteria (orange or blue) inside fluorescent phagosomes indicating the amoeba can ingest both the producer strain and the mutant. **(C)** Representative 2D maximum projections of tripartite co-cultures between amoeba and different ratios of wt and Δ*pys* cells. Orange and blue arrow heads indicate wt and Δ*pys* cells inside phagosomes respectively. **(D)** Comparison of the phagocytic indices in bipartite co-cultures of bacteria with amoebae. **(E)** Comparison of the phagocytic indices in tripartite co-cultures with different ratios of wt and Δ*pys* cells. Amoebae can ingest both the strains and does not exhibit any preferential feeding behaviour. Both **(D)** and **(E)** were performed in three independent replicates. Graphs are representative of three independent experiments. Data shown are mean ± standard error (n=10 images per ratio). Mann-Whitney test was performed to evaluate the statistical significance between the phagocytic indices of wt and Δ*pys* strains. Inter-ratio comparison (between 1:3 and 3:1) was carried out using a two-way ANOVA followed by Holm-Šídák’s multiple-comparisons test. ns not significant. Scale bars are 10 μm. The following source data and figure supplement are available for figure 2: **Source data 2.** Related to Figure 2D. **Source data 3.** Related to Figure 2E. **Figure supplement 1.** Comparison of the phagocytic indices among the co-culture.

In the tripartite co-culture with a 1:1 bacterial mixture, the PIs of both bacteria remained comparable (Figure 2E). The inter-ratio comparison among tripartite co-cultures (with 1:3 and 3:1 bacterial mixtures) revealed that the PI of wt in the 1:3 ratio was comparable to the PI of Δ*pys* in the 3:1 ratio. A similar trend could be observed between the PI of Δ*pys* in the 1:3 ratio and the PI of wt in the 3:1 ratio (Figure 2E). The variation in the PIs of both wt and Δ*pys* in the 1:3 ratio could be reversed in the co-culture with a 3:1 ratio of the bacteria (Figure 2—figure supplement 1). Overall, these findings show that amoebae do not exhibit any feeding preference towards the producer or the mutant strain.

### Alterations in nutrient availability leads to diminished production of pyreudiones

Since *P. fluorescens* wt was found to be ingested by amoebae, the role of pyreudione A production on the temporal dynamics of this amoebae−bacteria interaction was investigated. We performed a time course analysis on the production of pyreudione A in co-cultures. The co-cultures were prepared in two media. First, in a nutrient-rich medium, Peptone Yeast Glucose (hereafter PYG100) and second in a nutrient-poor medium, 10% of PYG100 medium (hereafter PYG10). Despite being a nutrient-poor medium, both amoebae and bacteria could still grow in PYG10 (Figure 4—figure supplement 1, Figure 4—figure supplement 2). The production of pyreudione A also occurred at lower titers in PYG10 when compared to PYG100 media (Figure 3—figure supplement 1). Since bacterial virulence and other evasion mechanisms have been reported to be mediated by cell density and other population dependant processes (Darch et al., 2012; Matz et al., 2008; Matz and Kjelleberg, 2005), We inoculated the bacteria with *D. discoideum* at a multiplicity of infection (MOI) of 5 or 100 bacteria per amoeba.

**Figure 3.**
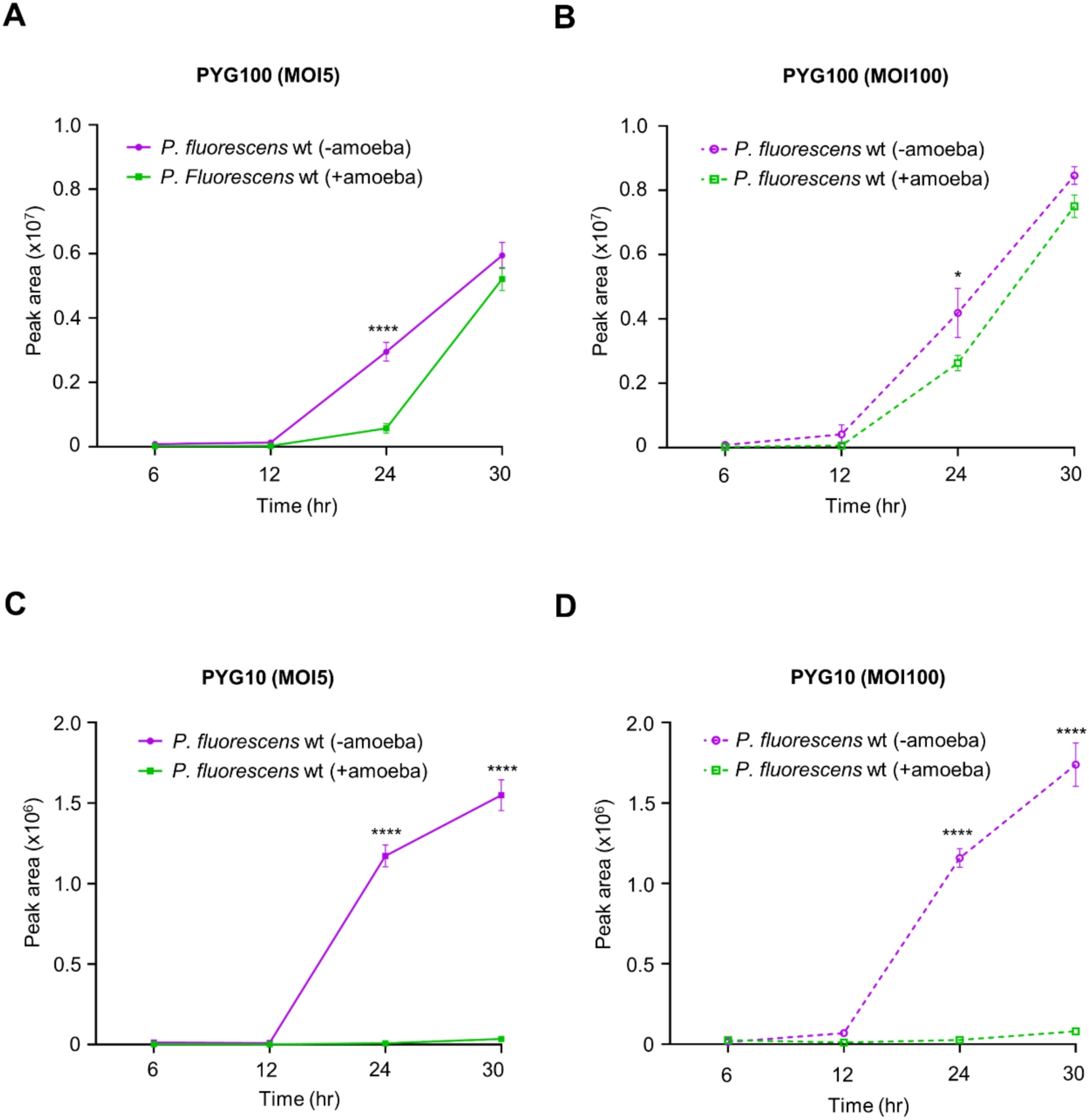
Production of pyreudione is diminished in co-cultures with nutrient-depleted media. **(A)** The production of pyreudione in the co-culture prepared with PYG100 media and bacterial population at MOI 5. **(B)** The production of pyreudione A in the co-culture prepared with PYG100 media and bacterial population at MOI 100. The levels of pyreudione gradually increases over time in co-cultures prepared with PYG100 media. **(C)** The production of pyreudione in the co-culture prepared with PYG10 media and bacterial population at MOI 5. **(D)** The production of pyreudione in the co-culture prepared with PYG10 media and bacterial population at MOI 100. Significant reduction in the production of pyreudione can be observed over time in the co-cultures prepared with PYG10 media. The production of pyreudione at each time point is represented as peak area of pyreudione A (at λ = 190 nm). Data shown are mean ± standard error pooled from three independent experiments. A two-way ANOVA with Holm-Šídák’s multiple-comparisons test was applied at each time point to determine the statistical significance. *p<0.05, ****p<0.0001. The following source data and figure supplement are available for figure 3: **Source data 4.** Related to Figure 3A. **Source data 5.** Related to Figure 3B. **Source data 6.** Related to Figure 3C. **Source data 7.** Related to Figure 3D. **Figure supplement 1.** Production of pyreudione A by *P. fluorescens* wt in different media.

**Figure 4.**
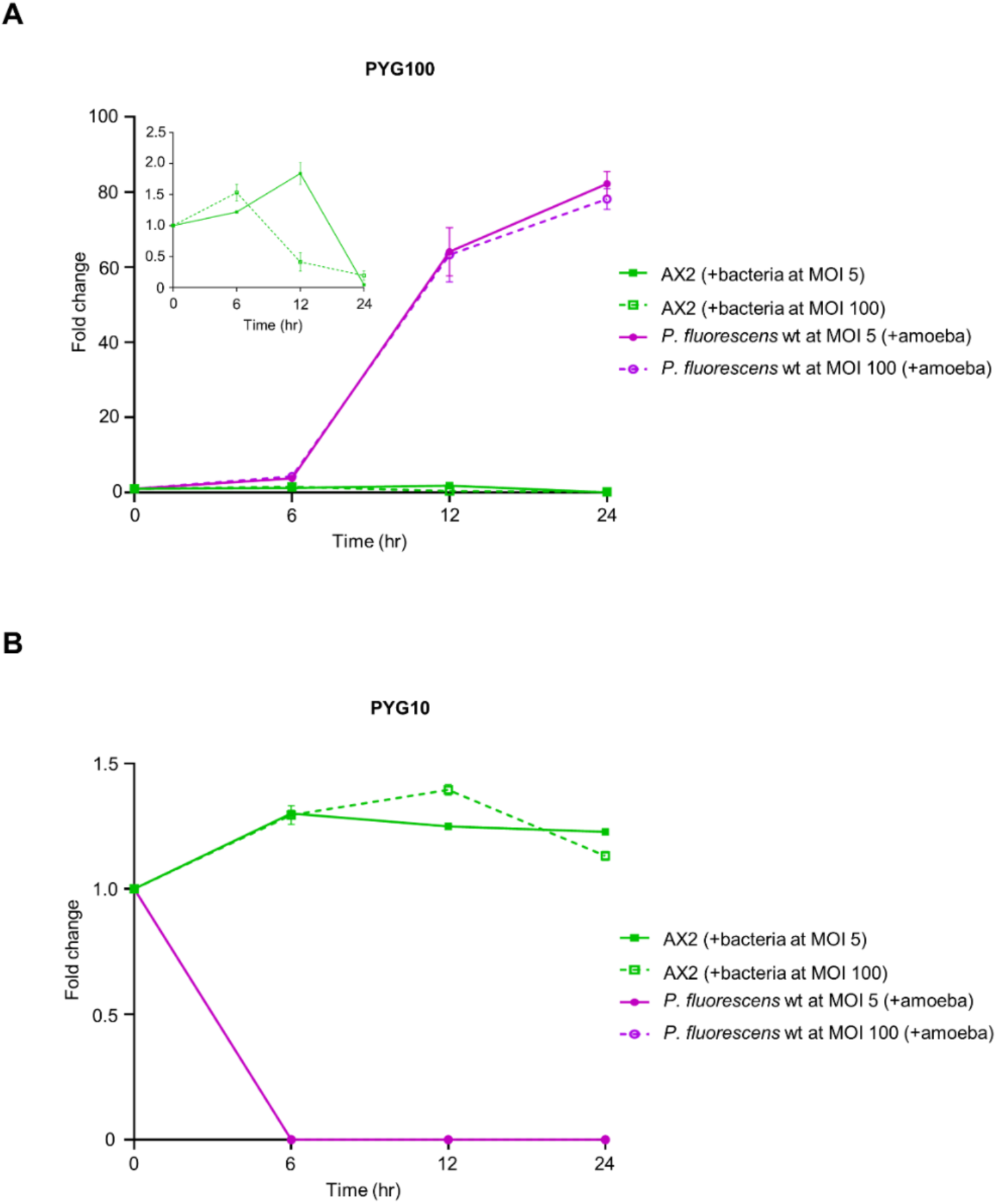
A strong Allee effect influences predator–prey relationship. **(A)** In PYG100 media, the bacteria population (at MOI 5 and 100) increases over time while there is a decline in the population of amoeba after 6 h. The differences in the growth of bacteria at both MOIs (5 and 100) were comparable. The inset shows an enlarged representation of the decline of amoebal population in co-culture over time**. (B)** In PYG10 media, the population of amoeba shows a slight increase but rather remain stable throughout the timepoints. But there is a significant decline in the bacterial population starting from 6 h. The differences in the growth of amoeba at both MOIs (5 and 100) were not significant. The change in amoebae or bacterial population is represented as fold change (y-axis) over time (x-axis). Data shown are mean ± standard error pooled from three independent experiments. Statistical significance was determined using two-way ANOVA followed by Holm-Šídák’s multiple-comparisons test at each time point. The following source data and figure supplement are available for figure 4: **Source data 8.** Related to Figure 4A. **Source data 9.** Related to Figure 4B. **Figure supplement 1.** Growth of *P. fluorescens* wt in amoeba-conditioned media (ACM). **Figure supplement 2.** Growth of AX2 in different media.

In the PYG100 co-culture, the production of pyreudione A gradually increased with time at both inoculation densities (MOI 5 and 100). At the final time point (30 h), the level of pyreudione A in the co-culture was comparable to the monoculture of *P. fluorescens* wt in PYG100 media (Figure 3A and B).

Interestingly, in the co-culture with PYG10, at both inoculation densities (MOI 5 and 100) the production of pyreudione A was significantly diminished when compared to the monoculture of *P. fluorescens* wt in the same media. The predation of bacteria from earlier time points could have caused the decline in the population of *P. fluorescens* wt and eventually led to diminished levels of pyreudione A (Figure 3C and D). This could also explain the lag in the production of pyreudiones in PYG100 media at an earlier time point when compared to the monoculture (Figure 3A).

Because PYG100 is a rich medium, bacteria outgrew the amoebae and increased the levels of pyreudione A in the co-culture. These findings demonstrated the influence of nutrition availability in altering the antipredator defence mechanisms.

### Allee effect influences microbial predator–prey relationship

To identify a potential link between cell density and the overall fitness of *P. fluorescens* HKI0770, we inoculated the bacteria with *D. discoideum* at two inoculation densities (MOI 5 and 100). We then determined the amoebal cell density and performed a CFU count assay of the bacteria. The time-dependent changes in the cell density of amoebae and bacteria were recorded from the co-cultures prepared in PYG100 and PYG10 media.

In the PYG100 co-culture, exponential growth of bacteria was observed at both inoculation densities (MOI 5 and 100) (Figure 4A). Even though an initial increase in amoebae cell numbers was seen, a decrease occurred as the bacterial population attained exponential growth (Figure 4A inset). Together with the quantifications of pyreudione titers within the co-cultures (Figure 3A and B), we concluded that bacterial growth in nutrient-rich PYG100 medium resulted in the production of pyreudiones leading to the decline of the amoebal population.

However, the co-culture in PYG10 medium, displayed amoebal cell numbers that were rather stable throughout the duration of the assay (Figure 4B). In contrast, there was a significant reduction in the bacterial population indicating that the amoeba was consuming bacteria (Figure 4B). At later time points, the bacterial population in co-culture was virtually depleted resulting in the absence of pyreudione A (Figure 3C and D). Additionally, we could also demonstrate that amoebal secretions did not alter the growth of bacteria (Figure 4—figure supplement 1), rather, predation by amoeba was the cause of the decline in bacterial population.

Alterations in the nutrition and variations in the bacterial cell density determined the level of pyreudione A in the system and therefore facilitated the predation of *P. fluorescens* HKI0770. Hence, in this predator–prey relationship, both bacteria and amoeba exhibit a direct link between population size and their individual fitness. We propose that this feedback mechanism is the result of a strong Allee effect. (Allee, 1949; Guin et al., 2023; Stephens et al., 1999)

### Interplay between nutrient availability and a strong Allee effect

A strong Allee effect is observed when the ability of a species to defend itself from the adverse effects of the other species diminishes with the population decreasing below a certain threshold.(Fadai et al., 2020; Stephens et al., 1999; Teixeira Alves and Hilker, 2017) Previous studies have shown the influence of Allee effects on amoebae−bacteria interactions.(DiSalvo et al., 2014; Rubin et al., 2019; Segota et al., 2022) However, the circumstances in which amoebae could overcome the potent evasion strategies of bacteria remain poorly understood.

To assess how nutrient availability coupled with a strong Allee effect determines the outcome of this predator–prey relationship, we analyzed the fruiting body formation of *D. discoideum* in co-culture with *P. fluorescens* wt. Three media (PYG100, PYG10 and Page’s Amoeba Saline Solution; PAS) with decreasing nutrient content and PAS being a non-nutrient saline (Tsai et al., 2023) were used. Different starting bacterial densities (MOI 5 and 100) were also tested.

In the wells with PYG100 media, the nutrient availability was sufficient for the bacteria to grow and compete with amoebal predation even at lower inoculation densities. This resulted in the production of pyreudiones and thereby killing the amoeba as described before.(Klapper et al., 2016) Having a high extracellular concentration of pyreudiones in a co-culture can effectively alter the feeding behaviour of amoeba (Figure 5—figure supplement 1). As a result, no fruiting bodies were observed on these wells (Figure 5).

**Figure 5.**
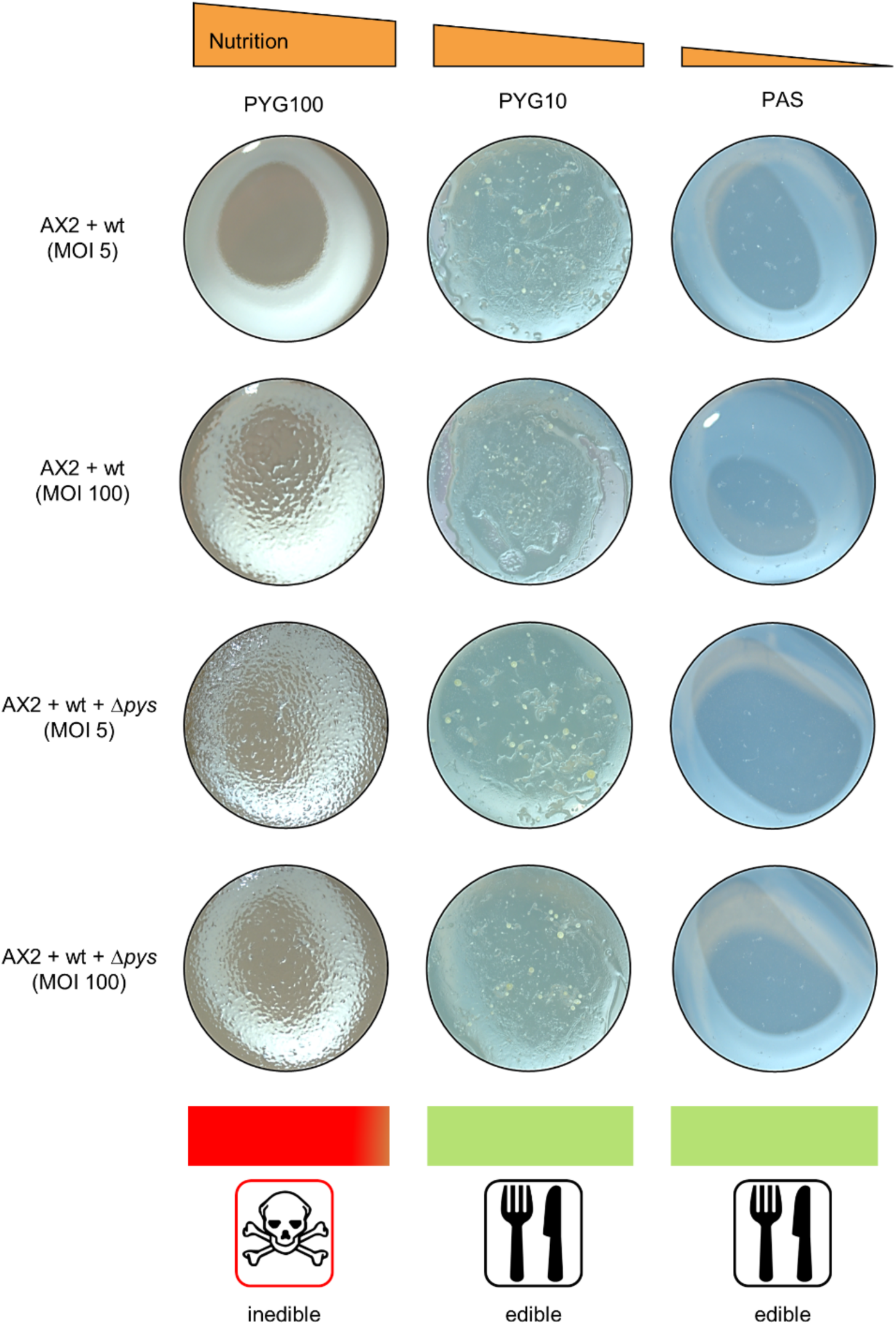
Interplay between nutrient availability and a strong Allee effect promotes the predation of *P. fluorescens* wt. Plaque assay with bipartite and tripartite co-cultures of amoeba and *P. fluorescens* HKI0770 strains. Fruiting bodies can be seen on the agar plugs as the nutrient availability decreases from left to right indicating predation by amoeba. Because PAS is a non-nutrient saline, the fruiting bodies on PAS agar are smaller than those in other media. The experiment was performed in two independent replicates. The following figure supplement is available for figure 5: **Figure supplement 1.** Co-culture of AX2 and Δ*pys* supplemented with *P. fluorescens* wt supernatant.

In the case of wells with nutrient-poor PYG10 and PAS, fruiting bodies of *D. discoideum* could be observed in the bipartite and tripartite co-cultures (Figure 5). In nutrient-poor media, bacterial growth did not reach exponential rates as the amoebae were able to feed on the bacteria and divide (Figure 5). The bacterial population did not reach the critical threshold required to produce enough pyreudiones to kill the amoebae (Figure 3C and D). Thus, creating a strong Allee effect on the bacteria which eventually leads to the decline of the bacterial population. With no bacteria remaining, amoebae started to aggregate and form fruiting bodies. Therefore, nutrient-depleted conditions and diminished pyreudione production provide a favourable environment for amoebal predation (Figure 3C and D, Figure 4B, Figure 5).

### Amoebal grazing alters the bacterial distribution

Predation of bacteria by protists (including amoebae) has been shown to cause changes in bacterial distribution and diversity.(Gao et al., 2019; Hahn and Höfle, 2001) To understand the interplay between amoebal predation and nutrient availability, we established a dual−agar system that allows monitoring predation while varying nutrient availability as encountered in many natural habitats. Two combinations of dual−agar systems (PYG100 – PYG10 and PYG10 – PAS) were prepared (See <u>Materials and</u> <u>methods</u>).

Co-cultures of amoeba and *P. fluorescens* wt were evenly distributed over these microcosms. In the PYG100 – PYG10 dual−agar system, bacterial colonies were more abundant on the nutrient-rich (PYG100) part of the microcosm without any signs of fruiting body formation (Figure 6A, Figure 6—figure supplement 1A). The remainder of the microcosm was, however, filled with fruiting bodies indicating bacterial consumption. This was further reflected by the distribution of pyreudione A in the microcosm. The nutrient-rich (PYG100) section of the microcosm had a higher concentration of pyreudione A when compared to the rest of the microcosm (Figure 6C). Bacterial proliferation is higher in the nutrient-rich part of the microcosm, leading to higher concentrations of pyreudione A resulting in the death of the amoeba. Lack of proper nutrition and predation pressure by amoeba resulted in the decline of the *P. fluorescens* wt population in the nutrient-poor section of the microcosm.

**Figure 6.**
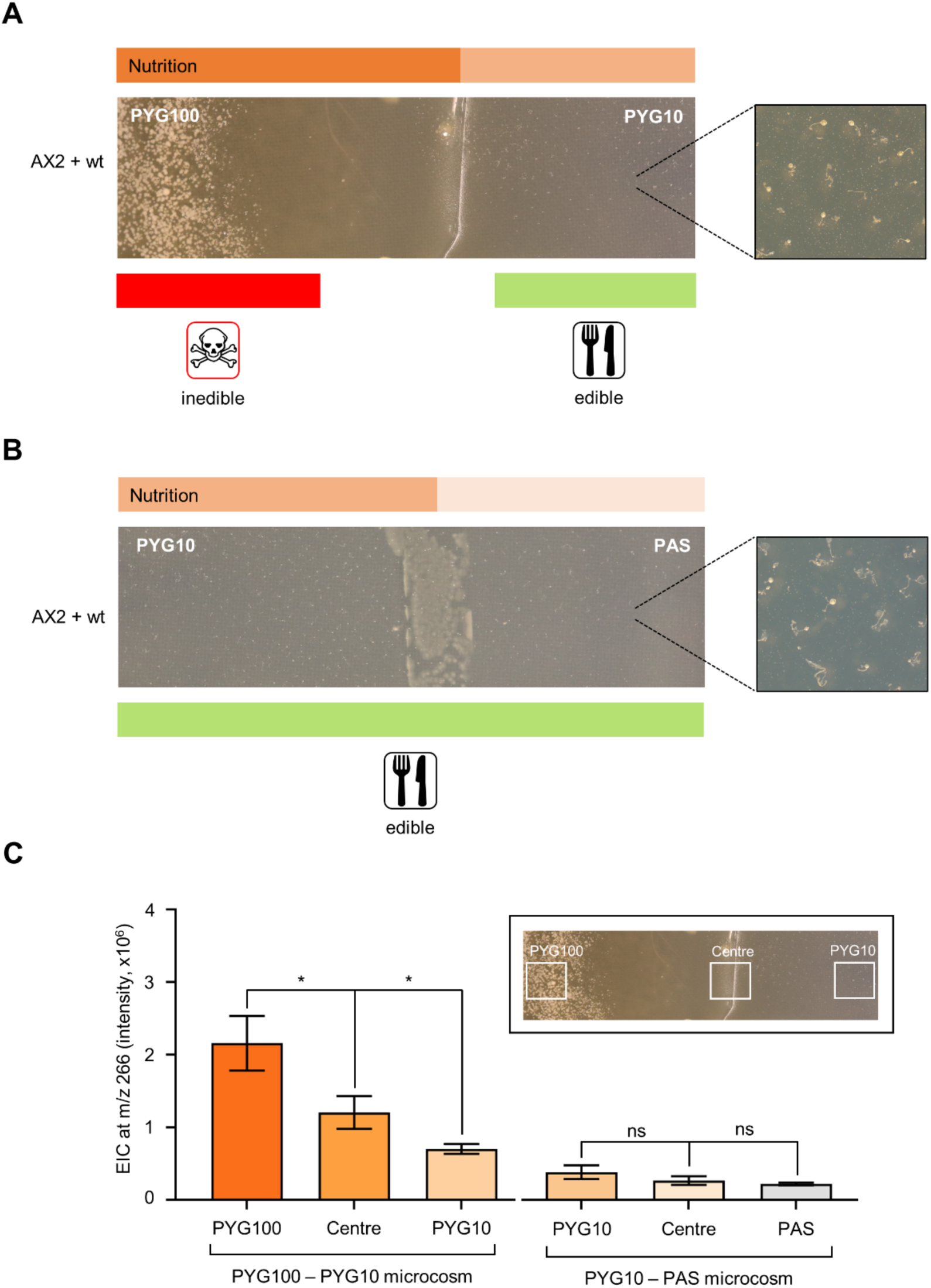
Co-culture of *D. discoideum* AX2 and *P. fluorescens* wt on dual–agar microcosm. **(A)** In the PYG100 – PYG10 microcosm, bacterial colonies can be seen towards the nutrient-rich (left) side of the microcosm. Fruiting bodies are spread throughout the rest of the microcosm. **(B)** In the PYG10 – PAS microcosm, fruiting bodies can be seen spread throughout both media. Insets show an enlarged view of the microcosm with the fruiting bodies. The experiment was performed in three independent replicates. **(C)** Comparison of the extracted-ion chromatogram (EIC) values of pyreudione A (m/z = 266) from different parts of the microcosms (PYG100 – PYG10 and PYG10 – PAS). The inset shows a representative image of the dual–agar microcosm with white squares indicating the areas from which agar pieces are obtained for UHPLC-MS. Data shown are mean ± standard error pooled from three independent experiments. An ordinary one-way ANOVA with Holm-Šídák’s multiple-comparisons test was applied to determine the statistical significance. *p<0.05. The following source data and figure supplement are available for figure 6: **Source data 9.** Related to Figure 6C. **Figure supplement 1.** Co-culture of *D. discoideum* AX2 and *P. fluorescens* wt::MRE-145 on dual–agar microcosm.

The microcosm composed of two nutrient-poor media (PYG10 and PAS) showed a different outcome. The presence of fruiting bodies across both media indicates that the bacteria were unable to produce enough pyreudione A to defend themselves against predation (Figure 6B, Figure 6—figure supplement 1B). The abundance of pyreudione A in different parts across the PYG10 – PAS microcosm was comparable (Figure 6C). Even though bacteria secrete low amounts of pyreudione A in PYG10 media (Figure 3—figure supplement 1), the lack of proper nutrition and a strong Allee effect generated by amoebal predation resulted in the decline of the *P. fluorescens* wt. Hence, our model emphasizes the critical role of nutrient availability and cell density in shaping predator−prey relationships.

## Discussion

Bacteria of the genus *Pseudomonas*, in particular *Pseudomonas fluorescens*, are ubiquitous soil microbes, well adapted to colonize the rhizosphere, promoting plant growth and acting as biocontrol agents. (David et al., 2018; Haas and Défago, 2005) However, the uncontrolled proliferation of bacteria has several adverse effects on the microbiome equilibrium. The predation of bacteria by protists, such as amoebae, in particular, maintains stable microbiome dynamics and functionality.(Gao et al., 2019) Conversely, predation exerts strong selection pressure on bacteria, which have evolved a wide range of antipredator strategies.(de Boer et al., 2019; Greub and Raoult, 2004; Matz and Kjelleberg, 2005; Queck et al., 2006; Zhang et al., 2021).

A notable antipredator strategy relies on the production of amoebicidal natural products – pyreudiones – by the soil-dwelling bacterium *P. fluorescens* HKI0770 (Götze et al., 2017; Klapper et al., 2016). This ability effectively turns the bacterium into an extracellular pathogen to amoebae. Due to its activity as a protonophore, the secreted pyreudiones induce intralysosomal acidification in amoebae thereby killing the predator. (Klapper et al., 2019).

Discriminating *P. fluorescens* HKI0770 wildtype and the Δ*pys* mutant during imaging were met with technical difficulties. Furthermore, the use of antibiotics as a selection pressure was not suitable during co-culture experiments with amoebae. To address this problem, different fluorescent tags were chromosomally inserted into the bacteria facilitating their differential identification. The chromatic bacteria toolbox is a transposon-based method used for differential fluorescent labelling of bacteria. (Schlechter et al., 2018) However, the use of this toolbox for intraspecific differentiation in intricate symbiotic relationships like of predator−prey interaction is a novel strategy. This also paves the way for potential applications in microbial identification and host−microbe interactions. By differentially labelling *P. fluorescens* HKI0770 wildtype and the Δ*pys* mutant, we investigated the amoebae−bacteria dynamics based on the amoebicide production.

The fluorescence imaging data reveal that *P. fluorescens* HKI0770 previously thought to evade predation,(Klapper et al., 2016) was, in fact, being ingested by the amoeba under certain conditions. This led us to investigate the circumstances under which amoebae are able to ingest the potentially amoebicidal bacteria. Our results also suggest that amoebae are not able to discriminate between the pyreudione producer strain and the Δ*pys* mutant as reflected by the similar phagocytic indices. Even though amoebae are able to sense and discriminate between edible and pathogenic bacteria, these patterns are mainly species-specific.(Lamrabet et al., 2020; Rashidi and Ostrowski, 2019; Schulz-Bohm et al., 2017) The inability of amoeba to perform intraspecific differentiation could explain the lack of a preferential feeding behaviour in amoeba.

Interestingly, the circumstances leading to the phagocytosis of the amoebicidal bacteria revolved around the availability of nutrition and cell density-dependent phenomena such as the Allee effect. Allee effects, in general, are population−dependant processes that are relevant for social and cooperative organisms. (Berec et al., 2007; Teixeira Alves and Hilker, 2017) A strong Allee effect arises when the growth rate of an organism declines below a certain threshold such that the same organism is unable to defend itself from the detrimental effects of another interacting organism. Eventually creating a negative impact on the per capita growth of the smaller population. (Fadai et al., 2020) Previous studies have shown that factors such as cell density and nutrient availability can alter bacterial virulence and thereby determine the outcome of predator−prey interactions.(Darch et al., 2012; DiSalvo et al., 2014; Haas and Keel, 2003; Sillo et al., 2011) Since pyreudione production is not mediated by quorum sensing, we thus hypothesised that the involvement of pleiotropic factors could influence the production of pyreudione A and thereby alter the feeding behaviour of amoebae. The time-dependent co-culture studies offered insights into the temporal dynamics of the amoebae−bacteria interaction. Notably, alterations in nutrient availability and a strong Allee effect within the co-culture system resulted in the diminished production of pyreudione A. Thereby rendering *P. fluorescens* HKI0770 vulnerable to predation by *D. discoideum*. We speculate that such a phenomenon likely mirrors ecological conditions found in natural habitats including the rhizosphere.(Gibson et al., 2011; Rounsaville et al., 2019)

Nutrient-rich hotspots in the soil such as the rhizosphere hold a vast repertoire of microbial biodiversity and a complex network of nutrient cycles.(Gao et al., 2019; Kuzyakov and Razavi, 2019; Sokol et al., 2022) This enhanced availability of nutrition promotes the soil-dwelling bacteria including *P. fluorescens* HKI0770 to multiply and secrete their antipredator natural products. As a result, bacterivorous organisms are unable to feed on the bacteria. As the bacteria move away from the rhizosphere, the nutritional availability and the microbial density tend to decrease.(Götze and Stallforth, 2020; Sokol et al., 2022) This results in the diminished secretion of toxic natural products. Thereby, rendering *P. fluorescens* HKI0770 vulnerable to free-living protists as proposed by our model (Figure 7).

**Figure 7.**
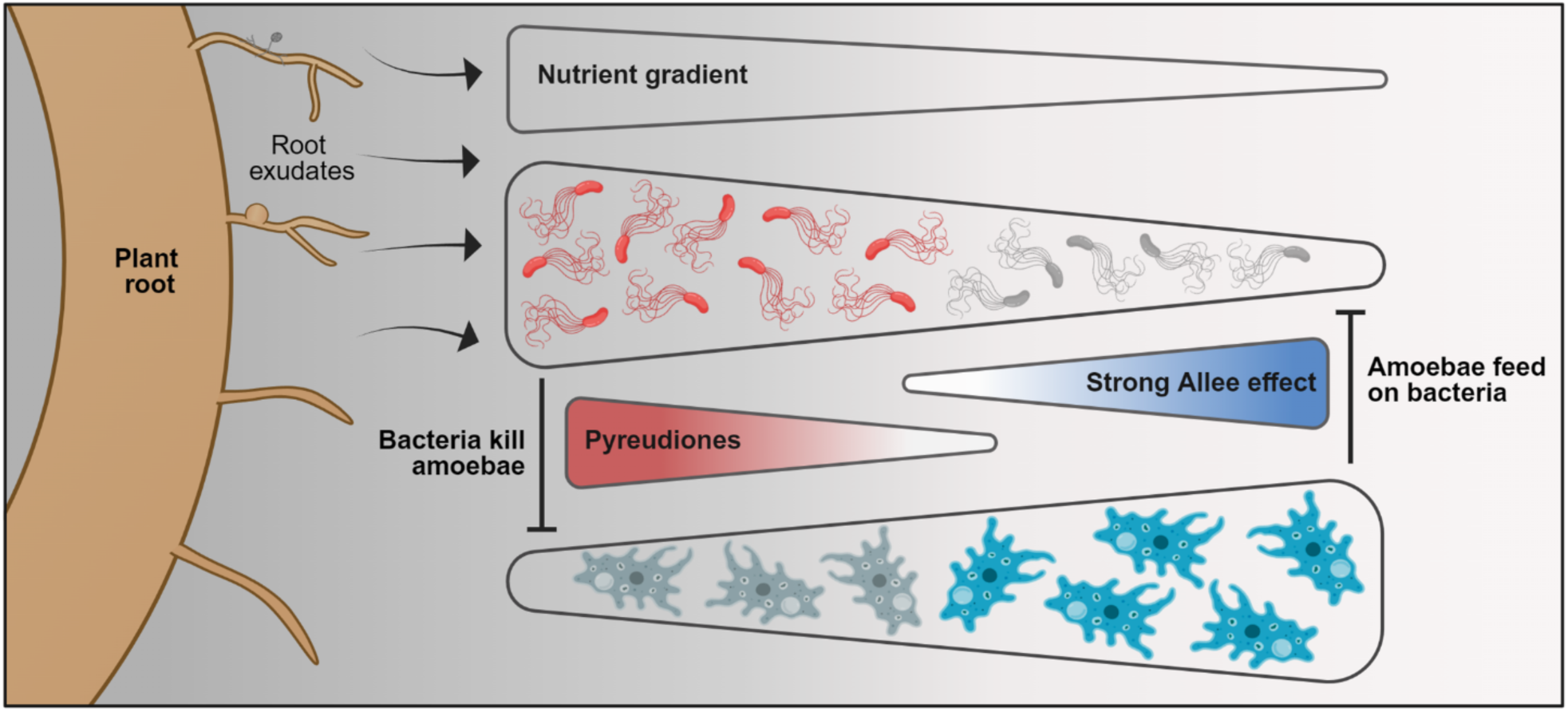
A schematic representation of the amoeba−bacteria interaction discussed in this study. The illustration was created with <u>BioRender.com</u>.

Our microcosm experiment showed that such phenomena could be replicated under laboratory conditions where the predator−prey dynamics are influenced by the Allee effect and various modes of nutrient distribution. We acknowledge that our microcosm experiment lacks several parameters including structural heterogeneity that often prevail in natural habitats. Still, our study distils the relevant underlying principles of the system and clearly demonstrates the multifaceted nature of predator−prey interactions and the influence of environmental conditioning in shaping them. These findings significantly contribute towards understanding the mechanisms underlying the predation-evasion strategies of bacteria and how amoebae can form various symbiotic associations with bacteria.

Amoebae like other protists, are top-down regulators for the microorganisms in the rhizosphere.(Gao et al., 2019) Their influence on the nutrient turnover of the rhizosphere in turn promotes plant growth. They also serve as a niche for pathogen evolution for both plant- and human-pathogenic bacteria.(Adiba et al., 2010) However, the importance of protists in microbial communities is often underestimated. Interestingly, our findings shed light on the roles of protists in driving the structure and function of microbial communities, with broader implications for microbiome engineering.

In summary, we have examined the hypothesis that within a polymicrobial environment, a stable community configuration can be reached. We have applied a fluorescence-based methodology that facilitates intraspecific differentiation of identical bacterial strains in eukaryote−prokaryote interactions. Extensive imaging of amoeba−bacteria co-cultures gave us insights into the intricate nature of predator−prey interactions. We also provide a direct experimental demonstration on the pleiotropic control of the production of natural products by bacteria can alter the feeding behaviour of amoeba, turning inedible bacteria into edible ones and *vice versa*. By itself, such non-genetic regulation is crucial as it governs the outcome of, not just predator−prey interactions but the whole spectrum of symbiosis. This prompts questions regarding the synergy between pleiotropic and genetic regulation in shaping microbial communities.

## Material and Methods

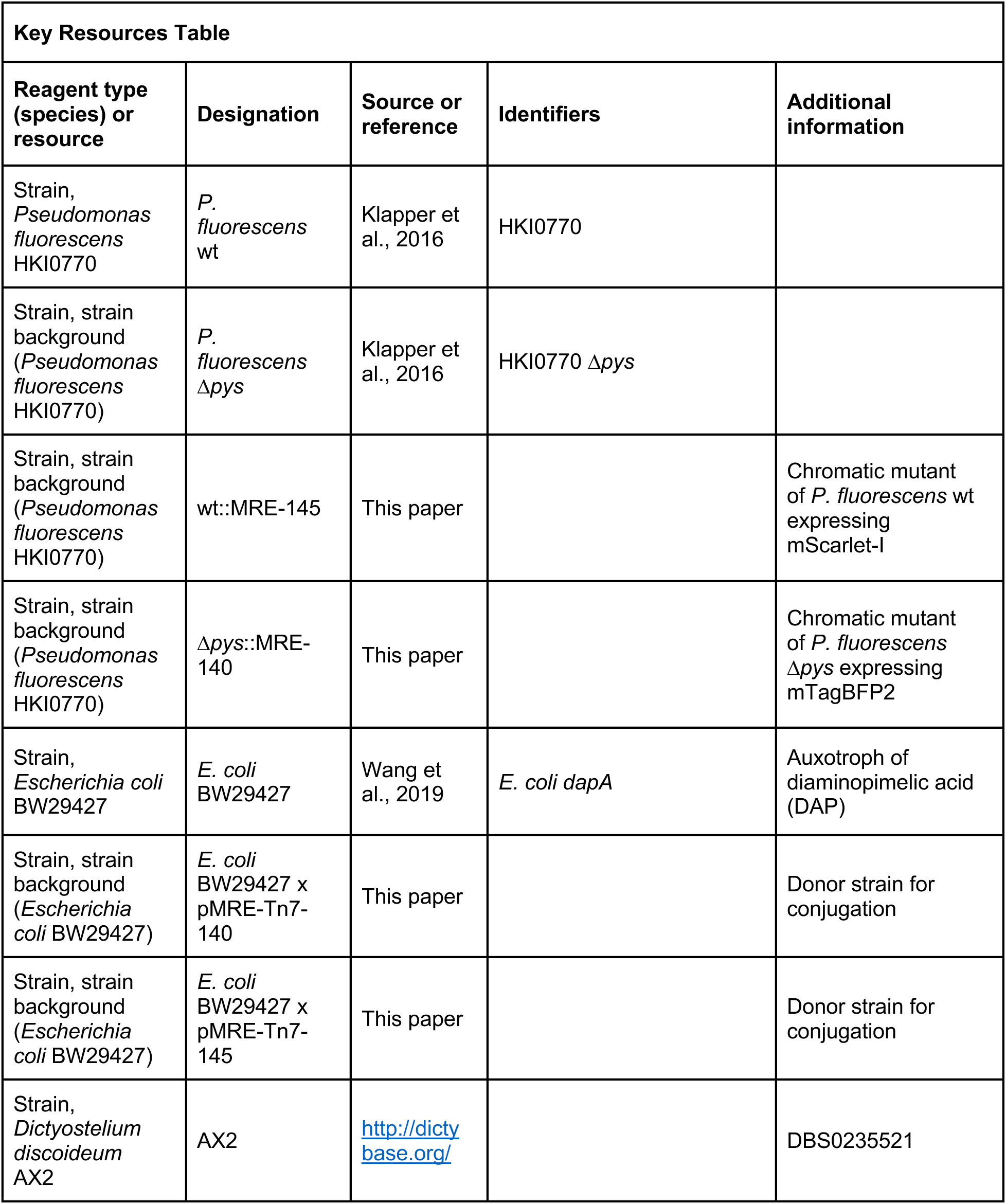
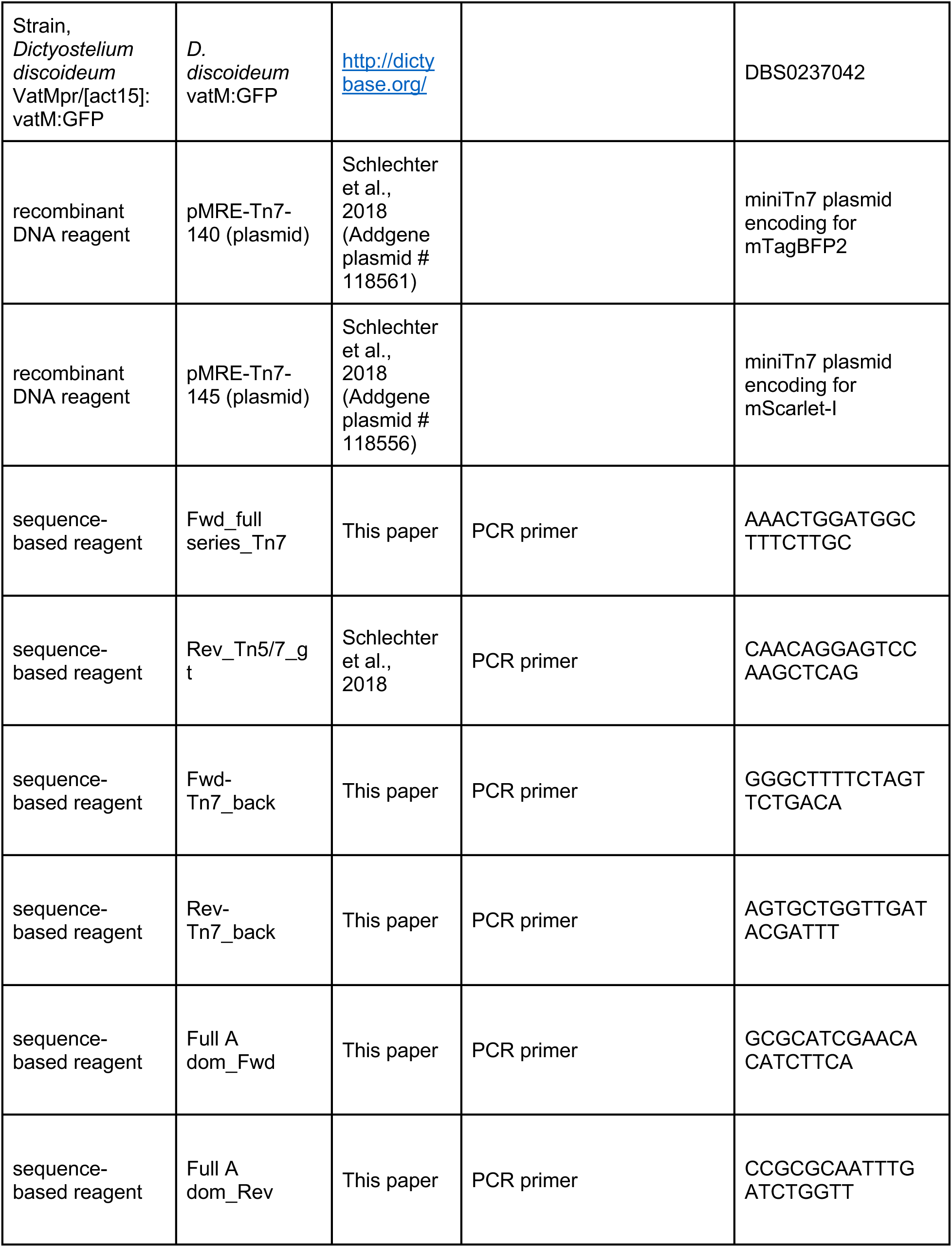
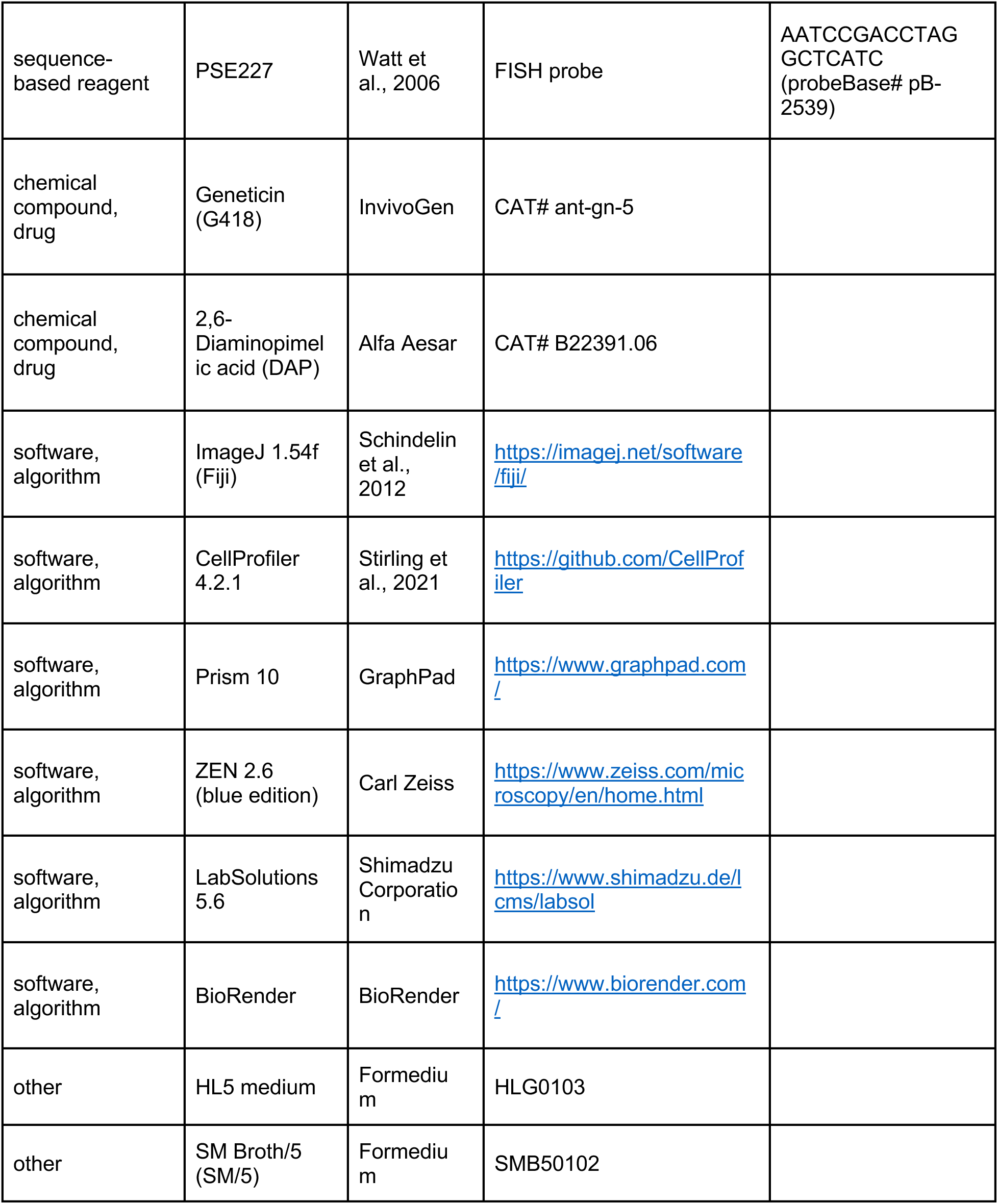

### Materials availability statement

Further information and requests and reagents should be directed to and will be fulfilled by the corresponding author, Pierre Stallforth.

#### Dictyostelium discoideum

*Dictyostelium discoideum* strain AX2 strain was grown in HL5 medium (Formedium^TM^, UK) supplemented with 1% (w/v) glucose (Carl Roth, Germany). *Dictyostelium discoideum* strain VatMpr/[act15]:vatM:GFP (hereafter *D. discoideum* vatM:GFP) strain was grown in HL5 medium supplemented with 1% (w/v) glucose and 50 μg mL^−1^ geneticin (G418, InvivoGen, France). Both strains were grown at 22°C in cell culture dishes or shaking flasks at 140 rpm.

#### Pseudomonas fluorescens HKI0770

All *Pseudomonas fluorescens* HKI0770 strains (wildtype (wt), Δ*pys* and chromatic mutants) were cultured at 28°C on a gyratory shaker at 180 rpm in SM/5 (Formedium^TM^, UK), unless mentioned otherwise. Before an experiment, the cells from overnight cultures were harvested by centrifugation (6000 x *g* for 5 min) and washed twice in 1x KK2 buffer (16.1 mM KH_2_PO_4_, 4 mM K_2_HPO_4_, pH 6.8).

#### Chromosomal insertion of fluorescent proteins

The chromatic bacteria toolbox (Schlechter et al., 2018) was implemented for the chromosomal integration of fluorescent tags into *P. fluorescens* wt and Δ*pys* strains. The plasmids, pMRE-Tn7-140 (encoding for mTagBFP2) and pMRE-Tn7-145 (encoding for mScarlet-I) were obtained from Prof. Dr Mitja Remus-Emsermann, Freie Universität Berlin, Germany.

Chemically competent *E. coli* BW29427 cells (obtained from Dr. Zhiying Zhao, DOE Joint Genome Institute, Berkeley, USA) were transformed with the pMRE-Tn7 plasmids via heat shock protocol (at 42°C for 30 s). The donor strains of *E. coli* BW29427 were grown in Luria-Bertani (LB) liquid medium at 28°C supplemented with 0.3 mM DAP (Alfa Aesar, USA) and 100 μg mL^−1^ ampicillin (Carl Roth, Germany). Conjugation with donor strains was carried out as previously described.(Klapper et al., 2023) The plasmids, pMRE-Tn7-145 and pMRE-Tn7-140 were delivered into *P. fluorescens* wt and Δ*pys* strains respectively. The donor and recipient cells from overnight cultures were harvested by centrifugation (6,000 x *g* for 3 min) and washed three times with LB liquid medium supplemented with 0.3 mM DAP. The recipient and donor strains were mixed in a ratio of 4:1 (donor:recipient). The conjugation mixes were drop-spotted on LB agar supplemented with 0.3 mM DAP and 0.1% w/v arabinose (Carl Roth, Germany) and grown at 28°C. The bacterial mix was resuspended on LB agar supplemented with 15 μg mL^−1^ gentamicin (Carl Roth, Germany) for selection of transformants. Chromosomal integration of fluorescent tags was confirmed by colony PCR (touchdown protocol) using Q5 HF Master mix (NEB, USA). The primer pair Fwd_full series_Tn7 with Rev_Tn5/7_gt were used to confirm the integration of fluorescent tags. The primer pair Fwd-Tn7_back with Rev-Tn7_back was used to confirm the absence of the delivery plasmid. The PCR cycling conditions were 95°C for 5 min, (95°C 30 s, 62°C 30 s; −1 °C/cycle, 72°C 2 min) × 15 cycles, (95°C 30 s, 58°C 30 s, 72°C 2 min) × 20 cycles, 72°C 2 min, and 8°C infinite.

#### Plaque assays with amoeba

The amoebicidal activity of *P. fluorescens* HKI0770 chromatic mutants was verified using a 24-well plaque assay as previously described.(Zhang et al., 2021) The assay was performed on SM/5 agar plugs present in each well of a 24-well plate (Sarstedt, Germany). Overnight cultures of *P. fluorescens* wt, Δ*pys* and their corresponding chromatic mutants were prepared as <u>described above</u>. To the individual wells, 30 μL of the respective bacteria was added and the plate was kept for drying.

The influence of the Allee effect (Allee, 1949) on amoebal predation was tested using another plaque assay. The assay was performed on Peptone Yeast Glucose (PYG100; 20 g L^−1^ proteose peptone, 18 g L^−1^ glucose, 2 g L^−1^ yeast extract in PAS), 10% of PYG medium (PYG10) and Page’s Amoeba Saline (PAS; ATCC medium 1323) agar plugs present in a 24-well plate. Cultures of *P. fluorescens* wt and Δ*pys* strains were prepared as <u>described above</u> and the OD_600_ was adjusted to 0.1. Bacterial cells were added to individual wells of each media at a multiplicity of infection (MOI) of 5 or 100 bacteria per amoeba. This initial bacterial inoculum was determined by colony forming unit (CFU) counts. Overnight culture of *Klebsiella aerogenes*, a typical food bacterium for *D. discoideum* was used as a positive control.

The influence of pyreudione A on amoebal predation was investigated by another plaque assay involving the supplementation of supernatant from *P. fluorescens* wt culture. The assay was performed on a 24-well plate with SM/5 agar plugs. Overnight cultures of *P. fluorescens* wt and Δ*pys* strains were prepared as <u>described above.</u> The supernatant from *P. fluorescens* wt culture was separated by centrifugation (6000 x *g* for 5 min) which was further made cell-free by passing through a 0.2 μm syringe filter (filtropur S plus, Sarstedt, Germany). Different dilutions of the supernatant were prepared in SM/5 broth (from 100% to 1%). To the individual wells, *P. fluorescens Δpys* was added at an MOI of 100 along with 50 μL of the respective dilution of the supernatant. The co-culture of AX2 and Δ*pys* strain without supernatant supplementation was used as a control.

For all the assays, *ca.* 50,000 AX2 cells were added to all the wells with bacterial lawns and air dried again. All the plates were incubated at 22°C until fruiting body formation occurs. Plaques generated as a result of fruiting body formation indicate that the bacteria are palatable and an absence of plaques indicate the amoebicidal phenotype.

The images were acquired using a stereo zoom microscope (Axio Zoom.V16, Carl Zeiss, Germany). The images were processed using ZEN 2.6 (Blue edition, Carl Zeiss, Germany) imaging software.

#### Detection of pyreudiones

The production of pyreudiones by the chromatic mutant as well as the production of pyreudiones by *P. fluorescens* wt in different media (PYG100, PYG10, and PAS) was analysed using Ultra-performance liquid chromatography−mass spectrometry (UHPLC−MS). Overnight cultures (5 mL) were extracted with 10 mL ethyl acetate. The organic phases were evaporated using a speedvac concentrator (ThermoFisher Scientific, USA). The crude extracts were dissolved in 200 μL methanol and filtered through a 0.2 μm PTFE syringe filter (Carl Roth, Germany). UHPLC analyses were carried out using the LCMS-2020 system (Shimadzu Corporation, Japan) equipped with a reverse-phase Kinetex^®^ C18 column (4.6 × 250 mm, 5 μm, 100 Å, Phenomenex^®^, USA). An autosampler was used to inject 10 μL of the methanolic extract. Using a flow rate of 0.7 mL min^−1^, peak separation was obtained with a linear gradient of acetonitrile in water with 0.1% (v/v) formic acid. The results were analyzed using LabSolutions 5.6 (Shimadzu Corporation, Japan) software. The total ion chromatograms (TIC) at the absorbance (λ) of 190 nm were extracted and stacked chromatograms were prepared by Prism 10 (GraphPad) software.

The production titre of pyreudione A in different media (PYG100, PYG10, and PAS) was also examined. Overnight culture of *P. fluorescens* wt prepared in Luria-Bertani (LB) liquid medium at 28°C on a gyratory shaker at 180 rpm was used to prepare 5 mL cultures in the three respective media with a starting OD_600_ of 0.5. These overnight cultures were further extracted using ethyl acetate and subjected to UHPLC−MS as mentioned above. The area under the curve (peak area) of pyreudione A, between the retention time of 5.7 and 6.0 min at λ = 190 nm was extracted for each media tested.

#### Differentiation of chromatic strains and Fluorescence *in situ* hybridization (FISH)

Overnight cultures of *P. fluorescens* HKI0770 chromatic mutants were prepared as <u>described above</u>. These were resuspended in SM/5 liquid medium and grown again to attain an OD_600_ of 0.5. The cells were harvested and washed twice with 1x KK2 buffer. The chromatic mutants were mixed in different ratios (1:1, 1:3, 3:1) and the mixtures were fixed using 4% paraformaldehyde (Sigma−Aldrich/Merck, Germany) for 4 h at 4°C.

FISH was performed using the probe, PSE227 (probeBase accession no. pB-2539) labelled with ATTO 465 dye (Eurofins Genomics, Germany). This probe targets the 16s rRNA region (227–245 bp) of *Pseudomonas* spp. The protocol performed was modified from (Zwirglmaier, 2010). A 10 μL aliquot from the various ratios was spread on a microscope slide with wells (Paul Marienfeld, Germany). After air drying, sequential ethanol dehydration (50%, 80% and 100% respectively) was performed. The slide was then incubated with hybridization buffer (1 pmol mL^−1^ PSE227 probe, 20 mM Tris-HCl, 0.9 M NaCl, 0.01% SDS, and 35% formamide, pH 7.5) at 46 °C for 2.5 hr. After hybridization, the slide was placed in pre-warmed wash buffer (20 mM Tris-HCl, 0.7 M NaCl, 5 mM Na_2_EDTA) for 20 min at 48 °C. The slide was then rinsed with ice-cold reverse osmosis water (RO; resistivity of 18.2 MΩ.cm at 25°C, stakpure, Germany). After drying, the cells were mounted in ProLong™ Glass antifade reagent (ThermoFisher Scientific, USA). A coverslip was placed over the slide and the edges were sealed using transparent nail polish.

#### Phagocytosis assay

Live-cell imaging was used to quantify the uptake of the *P. fluorescens* HKI0770 strains by amoebae. Before imaging, *ca.* 1 × 10^5^ *D. discoideum* vatM:GFP cells were seeded in an 8-well µ-Slide (ibidi, Germany) containing 300 μL of HL5 medium. After an hour of incubation, the media was replaced with 1x KK2 buffer. Cultures of the *P. fluorescens* HKI0770 chromatic mutants were prepared as <u>described above</u>. These were resuspended in 1x KK2 buffer to an OD_600_ of 0.1. Bipartite co-cultures of chromatic mutants with amoeba were prepared at a MOI of 300 bacteria per amoeba. Tripartite co-cultures were prepared using different ratios (1:1, 1:3 and 3:1) of the chromatic mutants with an overall MOI of 300. The slide was kept in the dark at 22°C for 30 min before imaging. The ingested fluorescent bacteria inside an amoebal phagosome were considered as “engulfed bacteria”. This was further represented as Phagocytic index (PI) (Chen et al., 2015; Sano et al., 2003) calculated according to the formula:

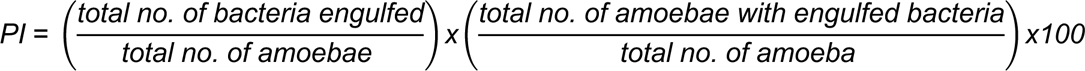

#### Fluorescence imaging

Imaging was performed using a spinning disk confocal laser scanning microscope (AxioObserver.Z1/7, Carl Zeiss, Germany) equipped with a 100x/1.40 NA Plan-Apochromat oil-immersion DIC M27 objective lens (Carl Zeiss, Germany). Images were captured using the diode lasers: 405 nm (50 mW), 445 nm (40 mW), 488 nm (50 mW) and 561 nm (50 mW) with the corresponding emission bandpass (BP) filters: BP 450/50, BP 485/30 and BP 629/62. For the <u>phagocytosis assay</u>, images were captured as z-stacks with 9–15 optical sections (1 µm per section) and 2 × 2 pixel binning.

#### Image analysis

After the acquisition, the images for <u>differentiation of the chromatic mutants</u> were subjected to deblurring (strength-0.3, blur radius-10 and sharpness-0.1) in the ZEN Blue software to improve their clarity. The images of bacteria from various ratios were imported into a CellProfiler (Carpenter et al., 2006) pipeline having three “RunCellpose” (Cellpose 1.0.2) modules to segment and count cells in three different channels. The pretrained “bact_fluor_omni” detection network with an expected object diameter of 15, flow threshold of standard 0.4 and a cell probability threshold of 0.2 was used. A total of 10 images from each ratio were used for the analyses.

The z-stack images from the <u>phagocytosis assay</u> were also subjected to deblurring (strength-0.4, blur radius-50 and sharpness-0.1) in the ZEN Blue software. 2D maximum projections of z-stacks were obtained using ImageJ 1.54f (Fiji) software. A total of 10 images from the bipartite and tripartite co-cultures were for the quantification.

#### Time−lapse detection of pyreudione in co-culture

Co-cultures of AX2 and *P. fluorescens* wt were set up in ø9 cm petri dishes (Sarstedt, Germany) with PYG100 and PYG10 liquid media. AX2 cells (*ca.* 2 × 10^6^) were seeded in the dishes and the bacteria was added at a MOI of 5 per amoeba and incubated at 22°C. As a control, monocultures of *P. fluorescens* wt were also prepared in both media. At the indicated time points (6 h, 12 h, 24 h and 30 h), the co-cultures (15 mL) were mixed thoroughly were extracted using 30 mL ethyl acetate (<u>detailed protocol</u> given above). After UHPLC, the peak area of pyreudione A (between the retention time of 5.7 and 6.0 min) at λ = 190 nm was extracted for each time point.

#### Fold change of amoebae and bacterial population

The co-cultures were prepared using PYG100 and PYG10 medium in ø9cm petri dishes as <u>mentioned above</u>. *P. fluorescens* wt was added to the co-culture at a MOI of 5 and 100 and the dishes were kept at 22°C. To determine the bacterial CFU count (CFU mL^−1^), the supernatant from the co-culture was collected and serial dilutions were carried out. A 100 μL aliquot from the final dilution was plated on LB agar and the resulting colonies were counted.

In order to obtain the AX2 cell numbers, 1x KK2 buffer was added to the petri dishes after the supernatant was removed. The adherent amoebae cells were scrapped out and mixed thoroughly. A 10 μL aliquot was added to a sample carrier slide (anvajo, Germany) and the cell numbers were determined by the fluidlab cell counter (R-300, anvajo, Germany).

Fold change is considered as a representative of fitness of a population.(Rubin et al., 2019) Hence, the fold change of the population (bacteria or amoeba) for each time point was calculated according to the formula:

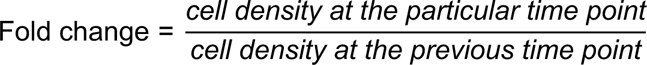

#### Dual–agar microcosm

To better understand how the Allee effect and nutrient availability affect the predator–prey relationship, a plaque assay was performed on a microcosm having two parts – one nutrient-rich and one nutrient-poor media.

To set up this system, 6 mL of PYG100 agar was poured into the wells of quadriPERM^®^ tissue culture dish (Sarstedt, Germany). Plastic inserts placed at the center of the wells helped to create two halves of the microcosm. Once that agar was solidified, the insert was removed and an equal volume of PYG10 agar was poured from the other side to create a dual–agar microcosm. A similar microcosm was also prepared using PYG10 and PAS media.

The microcosms were inoculated with AX2 cells (*ca.* 1 × 10^5^) along with *P. fluorescens* wt or the chromatic mutant at a MOI of 5. The amoeba–bacteria co-culture was spread throughout the wells using an inoculation loop. Once dried, the microcosms were incubated at 22°C. Images were acquired using a Canon EOS 800D camera and a stereo-zoom microscope when fruiting body formation occurs. Fluorescence images of the tissue culture dishes were captured using iBright™ CL1500 Imaging System (Invitrogen™, USA) equipped with a white Epi-LED source and a neutral density excitation filter (400– 700 nm).

In order to examine the distribution of pyreudione A in both microcosms, a 1 cm x 1 cm agar piece was obtained from the centre (overlap of both media) and from both sides of the microcosm (schematic shown in Figure 6C). The agar pieces were mixed thoroughly in ethyl acetate (10 mL) and subjected to sonication for 15 min. Extraction and UHPLC-MS was carried out according to <u>detailed protocol</u> given above. The extracted-ion chromatogram (EIC) of pyreudione A (m/z = 266) was from each part of the microcosm was extracted and used for comparison.

#### Growth of bacteria in amoeba-conditioned medium

Amoeba-conditioned media (ACM) was prepared using PYG100 and PYG10 media as described previously.(Brock et al., 2002) Briefly, AX2 cells (*ca.* 1 × 10^6^ cell mL^−1^) were inoculated in PYG100 or PYG10 media and grown under shaking conditions at 22°C for 18–20 h. After incubation, AX2 cells were harvested by centrifugation (500 x *g* for 10 min) and the ACM was further made cell-free by passing through a 0.2 μm syringe filter (filtropur S plus, Sarstedt, Germany). For the growth assay, 50% ACM (a 50/50 mix of the ACM and the respective culture media) was used.

Overnight cultures of *P. fluorescens* wt and Δ*pys* strains were prepared as <u>described</u> <u>above</u>. The OD_600_ of these cultures was adjusted to 1. A volume of 20 μL was transferred to a 96-well plate (Sarstedt, Germany) with a lid, followed by the addition of the respective ACM (180 μL) making a total volume of 200 μL. The OD_600_ measurements were acquired using a Tecan microplate reader (Infinite^®^ 200 PRO, Switzerland) every 15 min for 48 h. As a control, the strains were inoculated in normal PYG100 and PYG10 media.

#### Statistical Analysis

Prism (GraphPad) software was used to prepare the graphs and perform statistical analysis. Asterisks (*) are used in the figures to indicate different significance levels. To define significance, a p-value of less than 0.05 was used.

## Acknowledgement

We would like to thank Martin Klapper for providing the *P. fluorescens* HKI0770 strains. We thank Markus Günther and Rosa Herbst for providing insights throughout the course of this study and for providing comments on the earlier drafts of the manuscript. We thank Sebastian Pflanze, Lisa Reimer and Kevin Schlabach for their technical support and Sebastian Götze for the insightful discussions about natural products. We also thank Ruchira Mukerji for the initial support with FISH experiments. We acknowledge the use of Grammarly (https://app.grammarly.com) to improve the grammar and punctuation of the manuscript, but this tool was not used to generate any content in the manuscript.

## Additional information

### Author contributions

Conceptualization, H.R.S. and P.S.; Methodology, H.R.S and P.S.; Formal analysis, H.R.S.; Investigation, H.R.S.; Writing – Original Draft, H.R.S. and P.S.; Writing – Review & Editing, H.R.S. and P.S.; Funding Acquisition, P.S.; Supervision, P.S.

## Supplementary Information

**Figure 1—figure supplement 1.**
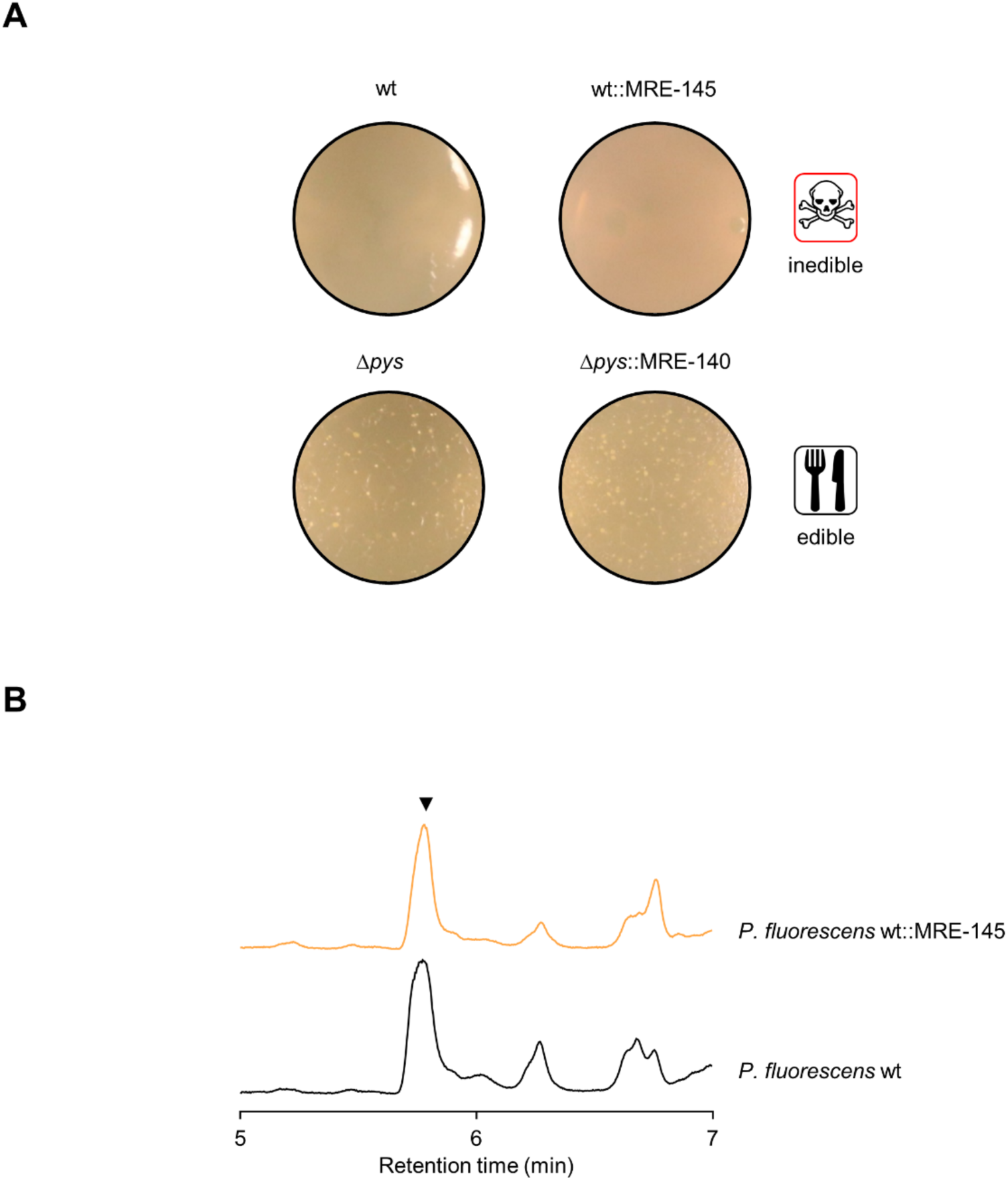
Pyreudione production of *P. fluorescens* HKI0770 chromatic strains. **(A)** Plaque assay with *Dictyostelium discoideum* AX2 and *P. fluorescens* HKI0770 strains. **(B)** Comparison of stacked chromatograms between the chromatic mutant and *P. fluorescens* wt. The phenotypes of the chromatic mutants remain identical to their parental strains. Black arrowhead indicates the peak for Pyreudione A (UV detection was at λ = 190 nm).

**Figure 2—figure supplement 1.**
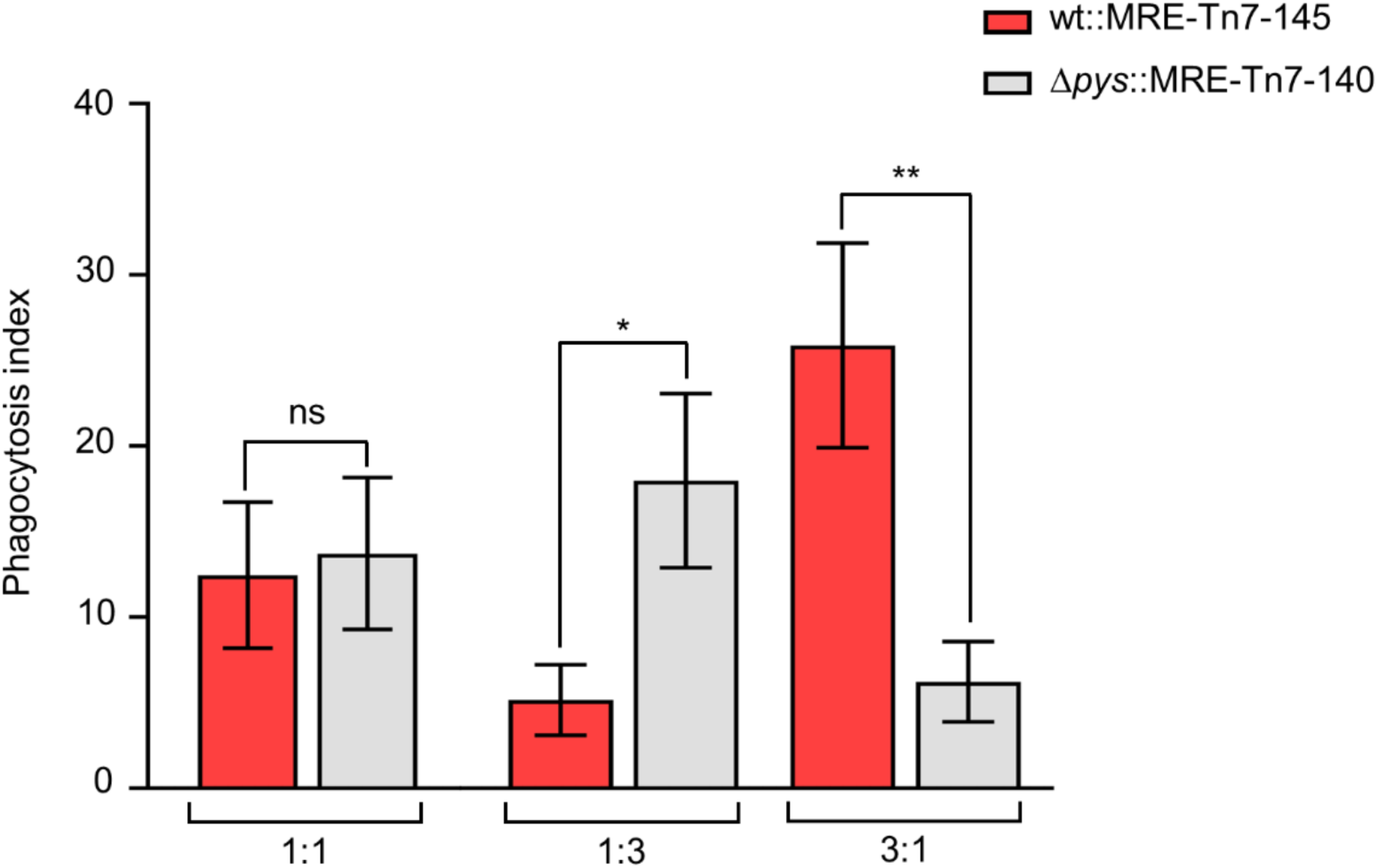
Comparison of the phagocytic indices among the co-culture. Quantification of phagocytic indices of wt and Δ*pys* strains in the different co-culture ratios (1:3, 3:1) based on the data shown in Figure 2E. Mann-Whitney test was performed to evaluate the statistical significance between the phagocytic indices of wt and Δ*pys* strains.

**Figure 3—figure supplement 1.**
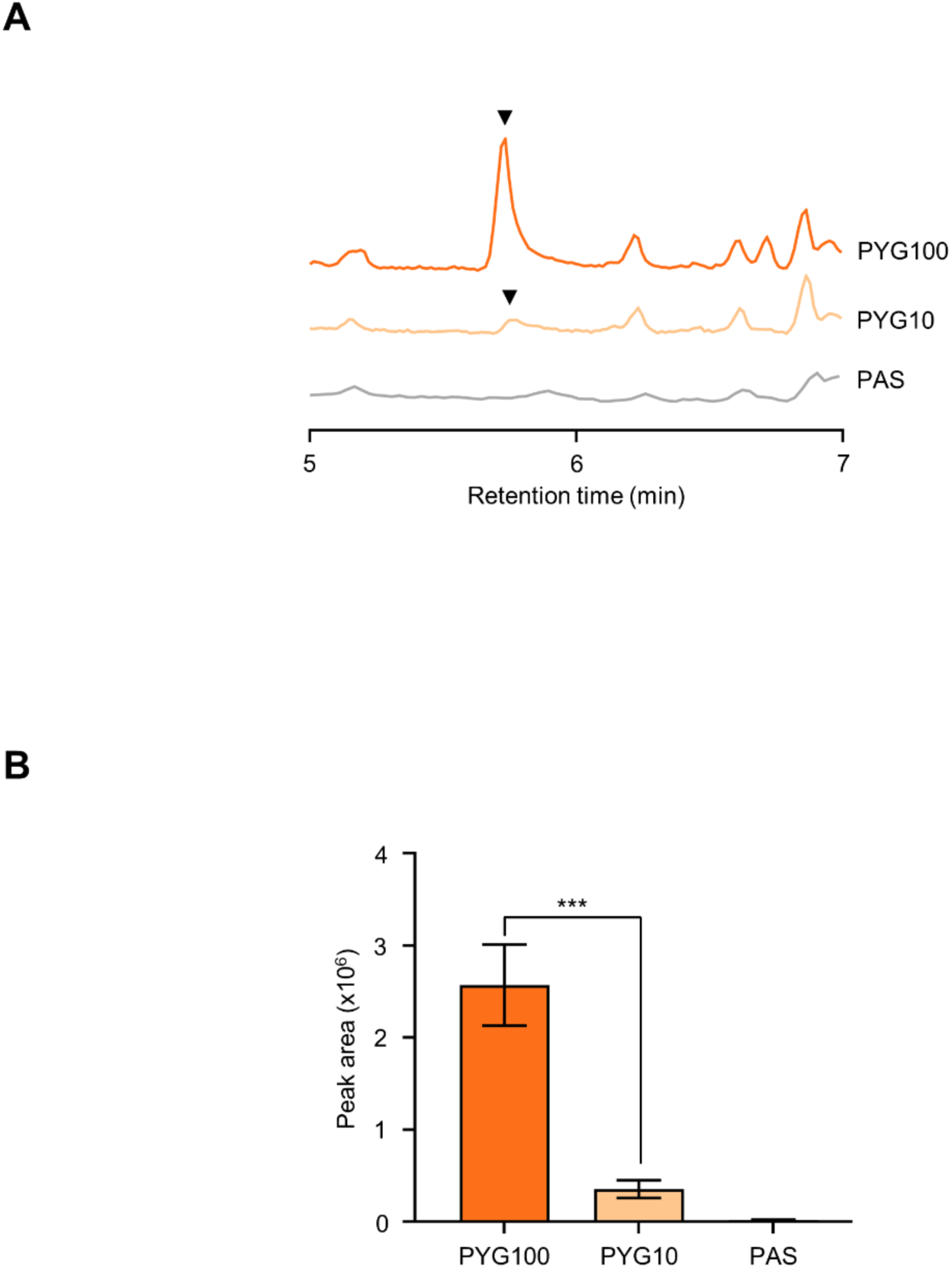
Production of pyreudione A by *P. fluorescens* wt in different media. **(A)** Stacked chromatograms showing the comparison pyreudione A production in PYG100, PYG10 and PAS. Black arrowheads indicate the peak for pyreudione A (UV detection at λ = 190 nm). **(B)** Production titre of pyreudione A in PYG100, PYG10 and PAS. Data shown are mean ± standard error (n=3 for each media). Ordinary one-way ANOVA with Holm-Šídák’s multiple-comparisons test was applied to determine the statistical significance. ***p<0.001.

**Figure 4—figure supplement 1.**
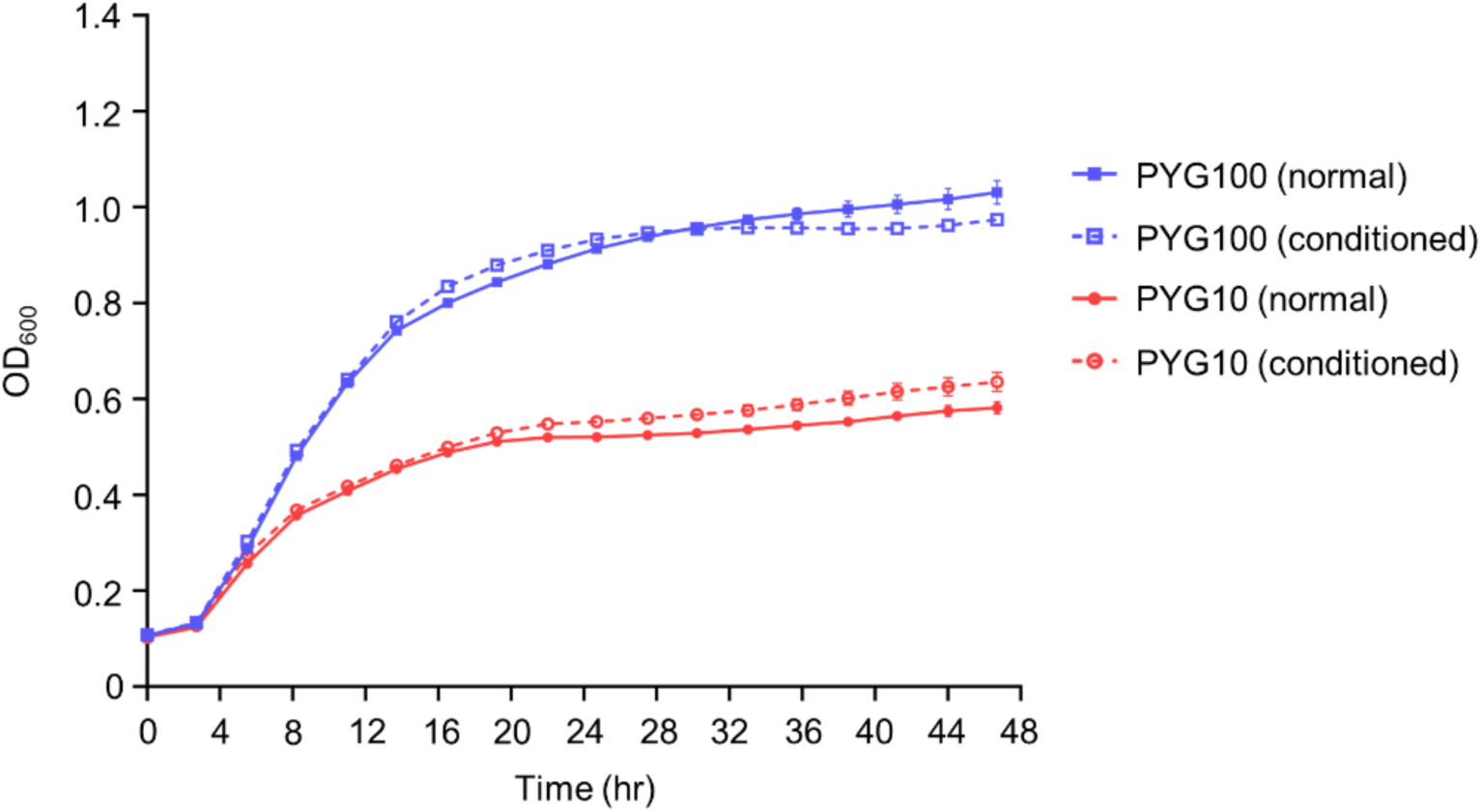
Growth of *P. fluorescens* wt in amoeba-conditioned media (ACM). The growth of *P. fluorescens* wt was comparable when grown in PYG100 and in PYG100 (ACM). A similar trend could be seen between PYG10 and PYG10 (ACM). Data shown are mean ± standard error (n=5 for each media). The experiment was performed in two biologically independent replicates.

**Figure 4—figure supplement 2.**
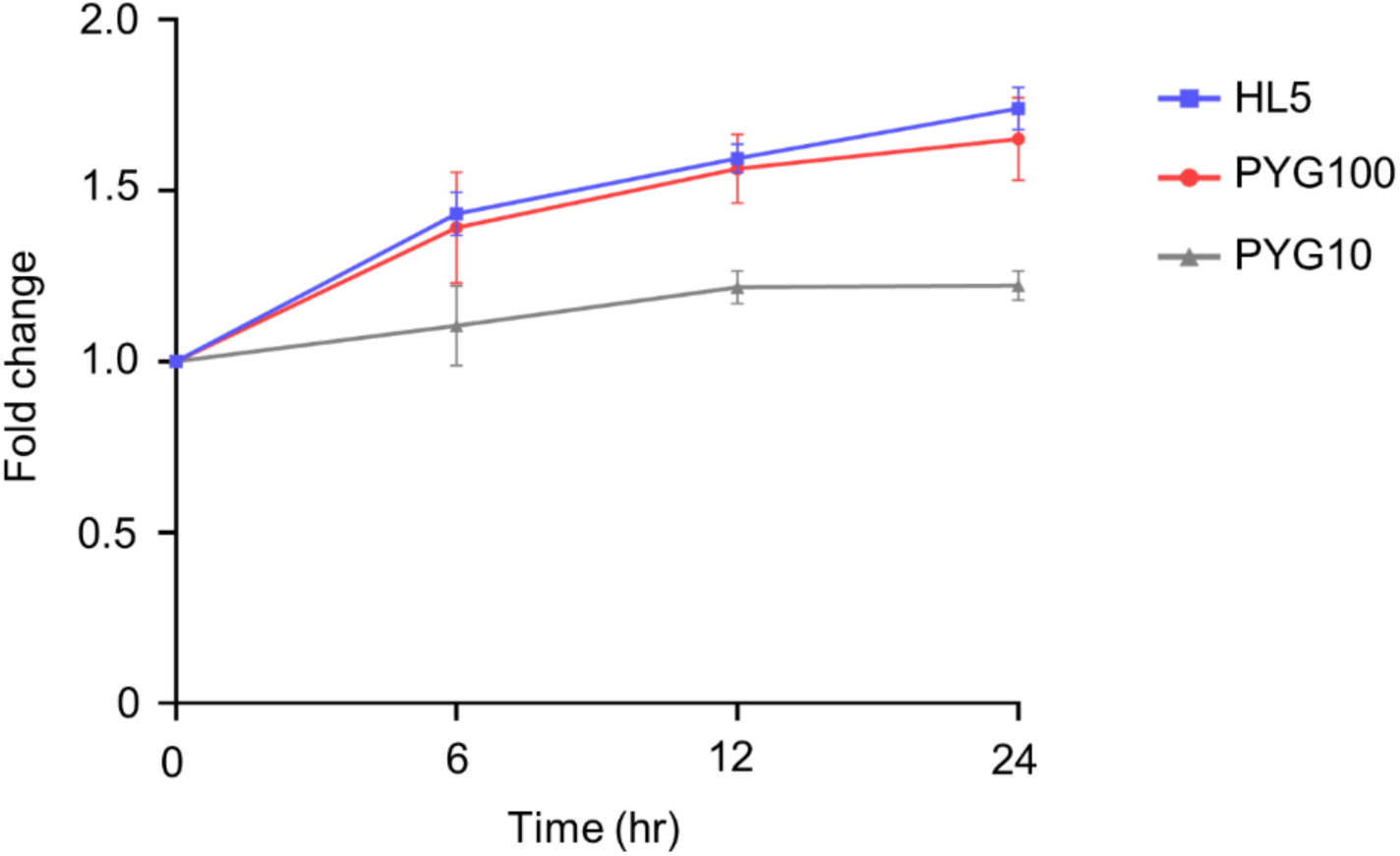
Growth of AX2 in different media. Axenic growth of *D. discoideum* AX2 in HL5, PYG100 and PYG10 media. The increase in cell density represented as fold change over time. Data shown are mean ± standard error pooled from three independent experiments. Statistical significance was determined using two-way ANOVA followed by Holm-Šídák’s multiple-comparisons test at each time point.

**Figure 5—figure supplement 1.**
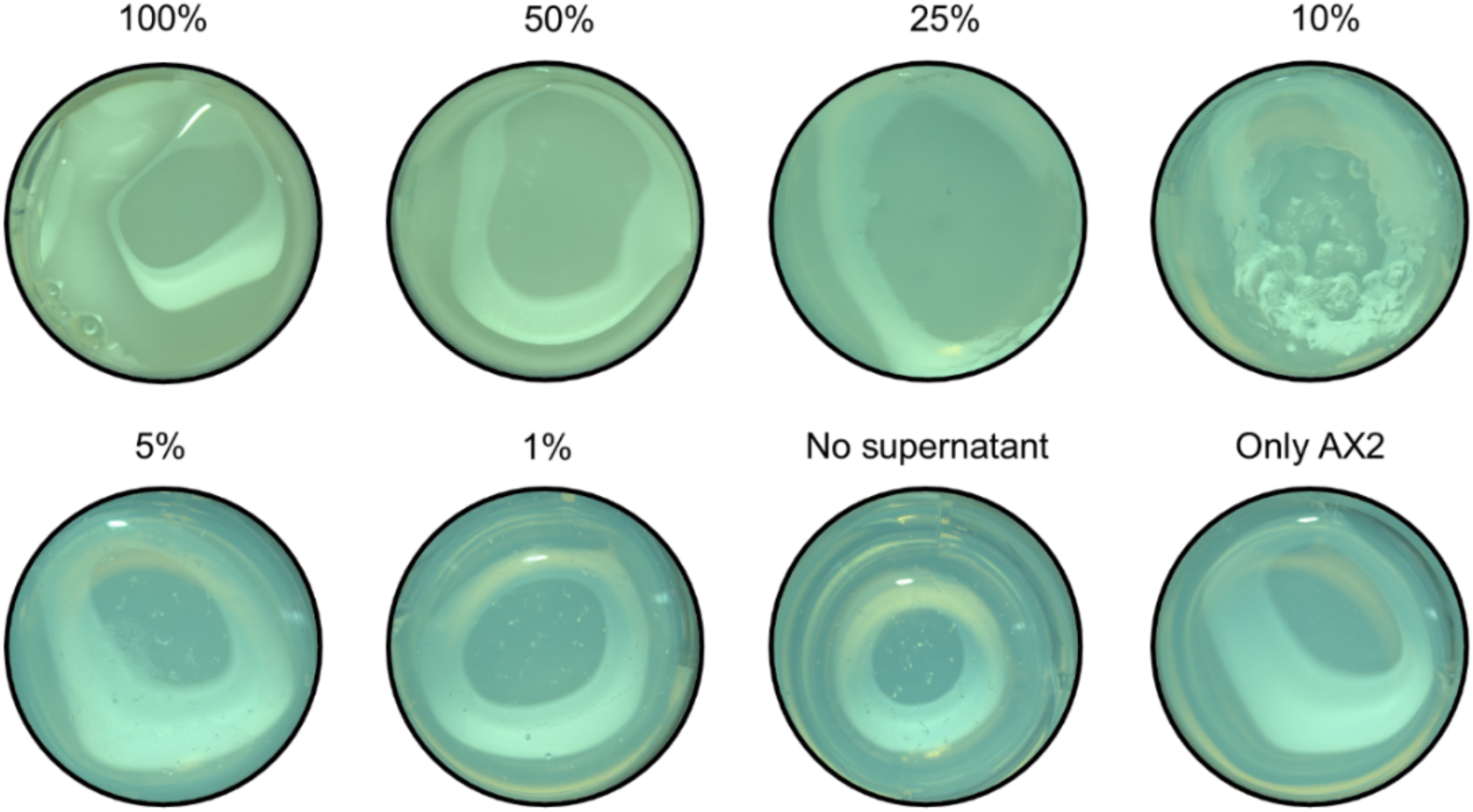
Co-culture of AX2 and Δ*pys* supplemented with *P. fluorescens* wt supernatant. Plaque assay with co-cultures of amoeba and *P. fluorescens Δpys*. As the concentration of *P. fluorescens* wt supernatant decreases from 100%, fruiting bodies can be seen on the agar plugs (5% and 1%) indicating influence of pyreudione A on amoebal predation. The experiment was performed in two independent replicates.

**Figure 6—figure supplement 1.**
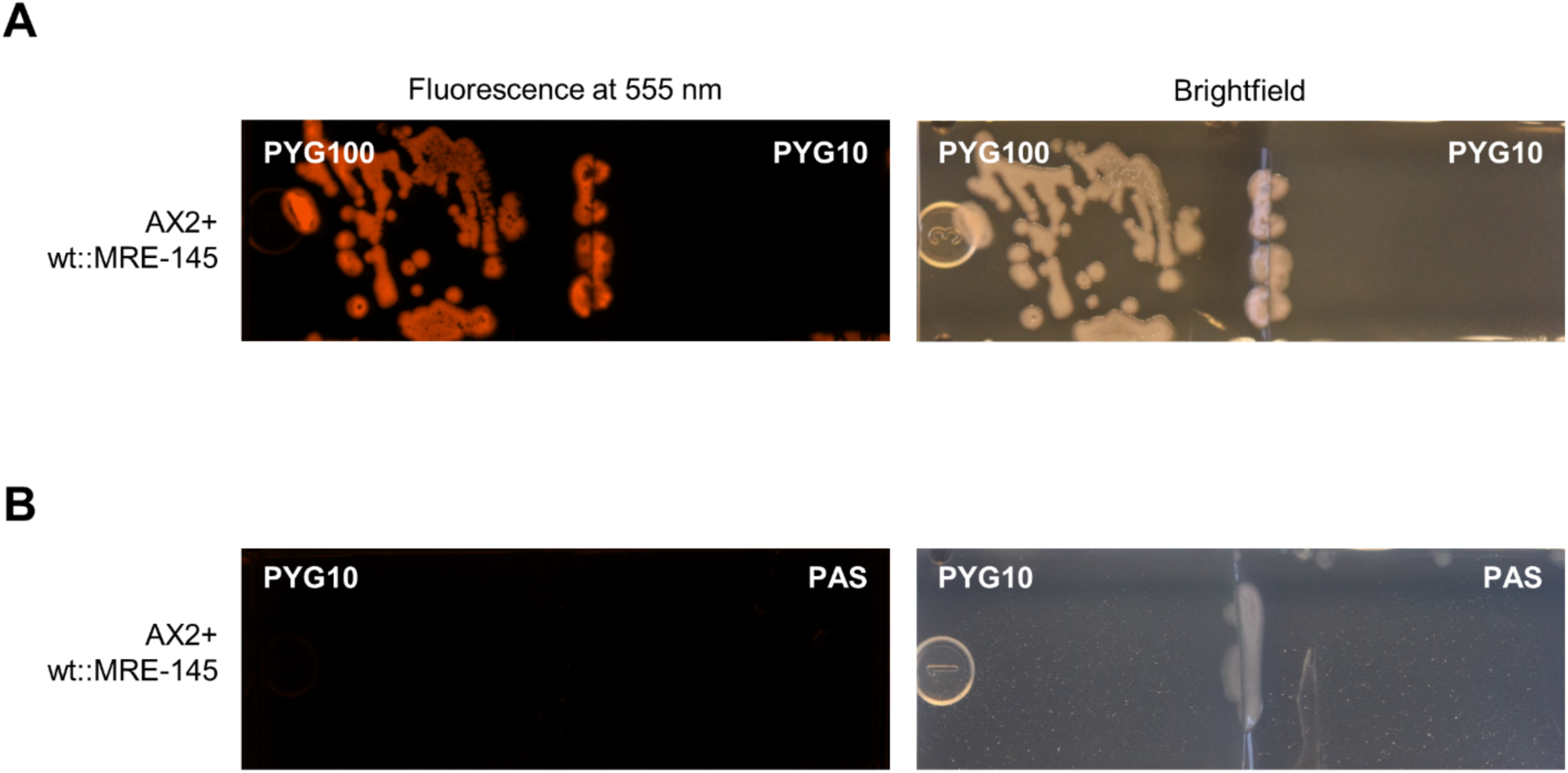
Co-culture of *D. discoideum* AX2 and *P. fluorescens* wt::MRE-145 on dual–agar microcosm. **(A)** In the PYG100 – PYG10 microcosm, fluorescent (orange) bacterial colonies can be seen distributed towards nutrient-rich (PYG100) left side of the microcosm. Absence of bacterial colonies on the rest of the microcosm indicate amoebal predation on the bacteria. **(B)** In the PYG10 – PAS microcosm, an absence of fluorescence and distribution of fruiting bodies can be seen throughout both media.

